# Conformational Dynamics in Insulin Receptor Kinase Reveals a Type III Allosteric Pocket

**DOI:** 10.1101/2025.05.10.653287

**Authors:** Jyoti Verma, Harish Vashisth

## Abstract

Allosteric modulation of kinases is a promising approach for pharmacological intervention, particularly for designing selective kinase modulators. However, the structural information on allosteric pockets remains limited for most kinases in the human kinome. In this work, we present comprehensive characterization of a type III allosteric pocket in the insulin receptor kinase (IRK) by uncovering its structural features and conformational dynamics. Specifically, we used microsecond-scale atomistic molecular dynamics (MD) simulations to investigate apo and inhibitor-bound IRK structures. Our findings suggest that the type III allosteric site is a “back pocket” in IRK, sandwiched between the N-terminal and the C-terminal lobes, beneath the *α*C-helix and the *β* - sheets of the N-terminal lobe. It has a hydrophobic cleft composed of both aliphatic and aromatic non-polar residues and a charge center to facilitate electrostatic interactions. We explored the binding of an experimentally known IRK inhibitor to the newly discovered allosteric pocket. Our results indicate that the *α*C-helix adopts an “out” conformation stabilized by the inhibitor which promotes an inactive conformation of the kinase. Furthermore, we observed a helical intermediate formation in the activation loop and a stable “DFG-out” conformation in both inhibitor-bound and unbound states. Our results also suggest that the residue M1051 in IRK functions as a gatekeeper residue, essential for maintaining the structural integrity of the *α*C-helix and regulating the binding of the allosteric inhibitor. Our findings are relevant for developing allosteric IRK modulators and informing therapeutic strategies targeting proteins in the insulin receptor family.

## 1 INTRODUCTION

Protein kinases are phosphotransferases crucial for activating various cell signaling cascades [1]. By transferring phosphate groups to substrate proteins, kinases regulate protein recruitment and function, effectively orchestrating almost all cellular processes [2, 3]. The human kinome consists of 518 protein kinases, of which 218 have been associated with human diseases [4, 5]. Hence, protein kinases have emerged as promising therapeutic targets for cancer and other metabolic disorders [6]. The discoveries of experimental structures of various kinases have aided in our understanding of their conserved structural elements, catalytic mechanisms, and peculiar features that differentiate kinase families and subtypes [7, 8, 9, 10]. The receptor tyrosine kinase (RTK) family constitutes multiple protein kinases evolved from a common ancestral gene that share a high degree of homology [11, 12]. For instance, the insulin receptor family of kinases consisting the insulin receptor (IR) and insulin-like growth factor 1 receptor (IGF1R) are RTKs, classified as protein homologs, which means they share significant structural and sequence similarities, despite having unique functions and ligand specificities [13]. Other such examples are Fibroblast growth factor receptor (FGFR), Vascular endothelial growth factor receptor (VEGFR) and Platlet derived growth factor receptor (PDGFR) [14]. Although some RTKs, such as EGFR and VEGFR, have been extensively studied, RTKs from the insulin family continue to be studied in terms of their structures and interaction sites (ligand binding pockets) [15, 16].

IRK and IGF1RK are multi-domain RTKs with a high structural similarity and overlapping downstream signaling pathways [17]. Each receptor consists of two extracellular ligand binding subunits (termed *α* -subunits) and two cytoplasmic subunits (termed *β* -subunits). Each *β* -subunit contains a large cytoplasmic region with inherent tyrosine kinase activity. Structurally, this tyrosine kinase domain comprises two lobes: a smaller N-terminal lobe featuring an *α* -helix (C-helix) and five *β* -sheets crucial for ATP binding, and a larger C-terminal lobe that facilitates phosphorylation and substrate binding [18]. The active site, responsible for ATP and substrate interaction, is situated between these lobes and includes the activation loop (A-loop) and the catalytic loop (C-loop). Due to the high homology between the kinase domains of IR and IGF1R, the development of specific inhibitors poses a challenge. Although IR plays a key role in regulating a variety of metabolic functions, IGF1R is mainly involved in growth regulation. Studies have reported the role of both IR and IGF1R in cancer development and progression [19]. More recently, it has been suggested that co-targeting IGF1R and IR increases the efficacy of cancer therapies [20, 21, 22, 23]. It has also been suggested that type 2 diabetes mellitus, and other conditions associated with insulin resistance like obesity and metabolic syndrome are significant risk factors for the progression of cancer [24, 25, 26, 27]. Thus, IR itself has emerged as a new target for cancer therapy; however, anti-cancer therapies specifically targeting IR and IR isoforms are still lacking [20].

Small molecule kinase inhibitors have become essential for treating a majority of diseases because of their favorable outcomes in clinical trials [28]. Kinase inhibitors are primarily categorized based on their binding site within the kinase domain. The type I and type II kinase inhibitors are small molecules that bind within the ATP binding pocket (orthosteric site) either to an active or inactive conformation of the kinase [29]. The type I inhibitors bind to the active conformation of the kinase, in which the A-loop is phosphorylated and is in the “DFG-in” conformation, and the type II inhibitors bind to an inactive kinase conformation, typically in a “DFG-out” conformation [30]. The type III and type IV inhibitors, on the other hand, are non-ATP-competitive; these are categorized as allosteric inhibitors that target sites distinct from the ATP pocket. The type III inhibitors bind between the small and large lobes adjacent to the ATP pocket, also known as the “back pocket” of the kinase [31, 32, 33]. Further, type IV inhibitors are those that bind to a site remote from the ATP binding pocket [34]. These inhibitors act as noncompetitive inhibitors, as their binding is unaffected by ATP. The majority of preclinically developed and clinically approved small-molecule inhibitors target the ATP-binding pocket of protein kinases [35]. However, achieving selective inhibition is challenging due to the extensive conservation and structural similarity of ATP-binding pockets. Targeting allosteric pockets has emerged as the new approach to address the issues of limited selectivity and drug resistance [36]. These allosteric sites, situated outside the highly conserved ATP-binding pocket, represent a promising alternative strategy for developing more selective and effective kinase inhibitors [37]. Although the number of identified allosteric inhibitors remains relatively limited compared to ATP-binding site inhibitors, the field of allosteric kinase inhibition has advanced rapidly in recent years [38, 39]. This progress is highlighted by the FDA approval of trametinib targeting mitogen-activated protein kinase kinase (MEK) [40], the clinical development of over 10 additional allosteric MEK and Akt inhibitors, and the first reports of allosteric inhibitors targeting other kinases [28].

Moreover, recent investigations have shown the prevalence of type III and type IV allosteric pockets in kinases [29]. A team at Novartis identified ten kinases with type III allosteric pockets and seven with type IV pockets [41, 29, 34]. The type III or “back pocket” ligands are associated with specific kinase conformations and proximity to the catalytic site [42]. Further analyses of kinase crystal structures have identified multiple co-cyrstallized allosteric ligands and reported 90 type III ligands across 19 kinases [43, 44, 45]. Based on these studies the type III pockets are typically characterized by: (i) their locations adjacent to the ATP binding pocket (orthosteric site), (ii) having an “out” conformation of the *α*C-helix, and (iii) a “DFG-out” conformation.

Despite significant focus on other kinase families, the insulin receptor family has received comparatively less attention regarding allosteric modulation. However, a series of allosteric modulators was identified as inhibitors of the insulin family of kinases through structure-activity relationship studies. Among these, a potent inhibitor MSC1609119A-1 was co-crystallized in complex with the kinase domain of IGF1R (PDB ID 3LW0) [46], and cellular assays confirmed its selectivity for IGF1RK over the homologous IRK. Moreover, we extensively investigated the structural basis for its selectivity by combining experimental structural data, small-molecule docking, and molecular dynamics (MD) simulations. This resulted in a comparative analysis of the binding of the allosteric inhibitor MSC1609119A-1 to IGF1RK and IRK, where we modeled the inhibitor interactions in IRK [47]. Our results highlighted the role of residue conformational dynamics in respective allosteric pockets of IGF1RK and IRK. Thus, our findings, which demonstrated the successful binding of an allosteric inhibitor to IRK, provided strong evidence for the existence of a type III allosteric binding pocket in IRK (Figure 1). This is particularly significant, given that the type III allosteric pocket in IGF1RK has been characterized with a co-crystallized allosteric inhibitor, thereby highlighting the significance of investigating this pocket in its homolog, IRK. However, the binding mode of the most potent IRK inhibitor in the allosteric inhibitor series [46] remains unknown. Hence, in this study, we characterized the type III allosteric pocket in IRK and studied conformational dynamics of structural motifs known to form this type III allosteric pocket. We used all-atom MD simulations of apo IRK and an allosteric inhibitor-bound IRK to rationalize key features associated with type III allosteric pocket and allosteric inhibitor interactions in IRK. Our findings provide valuable insights for the rational design of type III allosteric inhibitors targeting IRK and advance the development of highly specific allosteric modulators for the IRK family.

**FIGURE 1.**
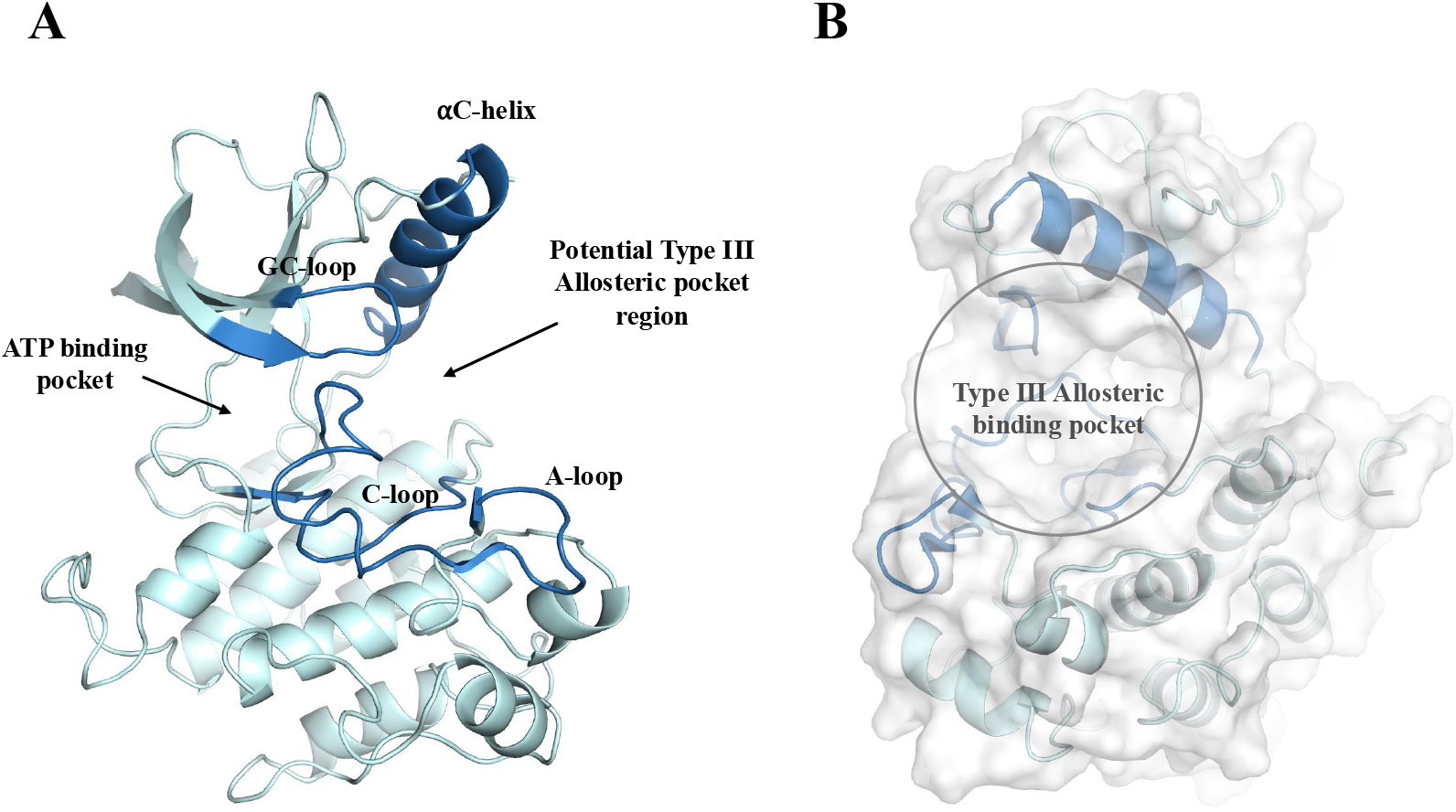
Potential allosteric pocket in the inactive state structure of IRK. (A) A cartoon representation of the inactive conformation of IRK (PDB ID: 1IRK). The *α*C-helix, GC-loop, catalytic loop (C-loop), and activation loop (A-loop) are highlighted in dark blue, while the remaining protein backbone is depicted in cyan. (B) A surface representation of the IRK structure showing the type III allosteric pocket region.

## 2 METHODS

### 2.1 System setup

To explore the existence of a type III allosteric pocket in IRK, we used the apo form of the IRK domain (PDB ID: 1IRK) as the starting structure [18]. We performed all-atom MD simulations of the apo IRK domain using the GROMACS-2020.4 package [48]. The protein atoms were parameterized using the AMBER ff99SB-ILDN force field [49]. A cubic simulation domain was created by placing the protein at the center with a minimum distance of 10 Å from the box edges. The TIP3P model for water molecules was used for solvation and Na^+^ and Cl^−^ ions were added to neutralize the net charge of the system. [50]. The final solvated and ionized system for apo IRK contained 56584 atoms. Following that, the simulation system was energy minimized for 50000 steps using the steepest descent algorithm [51]. After minimization, equilibration of the system was conducted in two steps, first in the NVT ensemble for 500 ps at 300K using the velocity rescale thermostat, followed by equilibration in the NPT ensemble for 10 ns using the Berendsen barostat at 1 atm pressure and 300 K temperature [52, 53]. Finally, the production MD simulations were carried out with an integration time step of 2 fs, and coordinates were saved every 100 ps. Three independent simulations, each 1 *µ*s (1000 ns) long, were conducted yielding a cumulative total of 3 *µ*s of simulation trajectory data.

### 2.2 Pocket exploration and characterization

The presence of the type III allosteric binding pocket in IRK was further explored by analyzing conformational ensembles based on MD simulation trajectories. We used MDpocket [54] to identify and characterize the binding pocket. The cavity detection in MDpocket is based on Voronoi tessellation, which partitions/subdivides the protein surface into grid-like regions (cells) based on the spatial arrangement of atoms. This geometric approach helps in identifying potential binding sites by focusing on spatial partitions that are likely to accommodate small molecules. The following steps outline the preparation (processing of the MD trajectory data), grid mapping, and analysis procedures used to examine the IRK structure.

#### 2.2.1 Preparation and Alignment

In the grid-based procedure used by MDpocket [54], the protein structure is mapped onto a grid to detect potential binding pockets. However, raw MD trajectory data often exhibit fluctuations or distortions due to periodic boundary conditions, which can introduce artifacts. To address this, each trajectory frame was aligned to the initial frame, which served as the reference structure. This alignment reduced the impact of artifacts by ensuring that each frame represented the conformational state of the protein relative to the reference structure (initial frame). This step ensures that any grid-based analysis performed subsequently is consistent and not influenced by distortions in the simulation data. After alignment, each frame was saved as an individual Protein Data Bank (PDB) file (total 10000 frames from each trajectory), facilitating systematic analysis and further processing. Solvent molecules and counter ions were removed before exporting the PDB files.

#### 2.2.2 Grid Mapping and Pocket Detection

To detect cavities, the protein structure was mapped onto a grid for each trajectory frame. A grid with a resolution of 0.5 Å was placed over the first snapshot of the aligned PDB files. Each grid point was associated with *α* -spheres, which are defined as spheres in contact with at least four protein atoms. These spheres were assigned to the grid point closest to their center. This grid point association process was iterated across all frames, using default settings that included a grid resolution of 0.5 Å, a minimum pocket size of 50 grid points, and a solvent-accessible surface area (SASA) model with a probe radius of 1.4 Å. The resulting data generated two key maps: a *density* map, reflecting the spatial distribution of *α* -spheres across the protein surface, and a *frequency* map, which indicates how often each grid point was occupied by an *α* -sphere across the trajectory. The density map was calculated by normalizing the number of *α* -spheres assigned to each grid point by the total number of frames. This provided insights into the spatial distribution of potential binding pockets, highlighting regions with higher concentrations of *α* -spheres. The frequency map was calculated by normalizing the number of grid point occupancies (where at least one *α* -sphere was assigned to a grid point) across all frames. The frequency map is useful to gain insights into the consistency and variability of potential binding sites. The grid points that were consistently occupied by *α* -spheres over the course of the simulation were indicative of more stable and accessible binding sites.

#### 2.2.3 Data Analysis

The pocket grid maps were further analyzed to identify and characterize the binding pocket. We identified the potential region of the type III binding site in the frequency grid map. For this selected pocket, the grid points with the isovalues 0.5 (i.e., grid points connected with *α* -spheres for at least half the simulation time) were retained to calculate pocket’s metrics (such as density, volume, etc.). The data from these maps offered insights into the variability and potential functional metrics to characterize the type III pocket in IRK. The grid maps were visualized and analyzed using VMD [55].

### 2.3 Docking and MD simulation of a known allosteric IRK inhibitor

A series of indole butyl amine derivatives have been shown to inhibit IRK activity experimentally [46]. However, the detailed structural characterization of the binding modes of these inhibitors remains elusive. In our previous study [47], we observed distinct differences in the binding mode of the inhibitor MSC1609119A-1 from this series when interacting with IGF1RK compared to IRK. In this study, however, we selected the compound showing the strongest inhibitory effect (lowest IC_50_ = 1.3 *µ*M) on IRK to illustrate its binding conformation within the predicted type III allosteric pocket of IRK.

We performed docking of the most potent allosteric inhibitor of IRK (N-1-[4-(5-cyano-1H-indol-3-yl)butyl]piperidin-4-yl-1H-indole-4-carboxamide) using AutoDock [56]. The inactive apo structure of IRK (PDB ID: 1IRK) was used as the receptor. Water molecules were removed, and hydrogen atoms were added to the protein structure to ensure correct protonation states. Subsequently, partial charges were assigned, and the receptor molecule was saved in PDBQT format, which contains the required atom types and charges for AutoDock calculations. Next, the three-dimensional structure of the inhibitor molecule was generated using MarvinSketch from ChemAxon. The ligand structure was converted to PDBQT format in AutoDock, which includes adding charges and specifying atom types. A grid box around the type III binding pocket region was generated with the x, y, and z dimensions of 20.609, 69.029, and 19.063, and these grid parameters were saved as the GPF configuration file. Next, the docking parameter file (DPF) was generated with 50 genetic algorithm runs, the population size set to 150, and the number of generations set to 27000. Finally, the docking simulation was executed using the prepared input files. The docked conformations were analyzed, and the most favorable binding pose with the lowest binding energy (most negative) was chosen for further MD simulations.

All-atom MD simulations were conducted on the IRK structure complexed with the allosteric inhibitor to investigate the conformational dynamics of the type III allosteric pocket as well as the inhibitor binding conformation. The IRK structure was parameterized using the AMBER ff99SB-ILDN force field while the inhibitor structure was parameterized with the general amber force field (GAFF) using the Antechamber module [57, 58, 59]. The coordinates were merged, and additionally, the inhibitor topology information was integrated into the system topology file. The system was solvated with TIP3P water molecules, and Na^+^ and Cl^−^ ions were added to neutralize the overall charge. The final solvated and ionized system for inhibitor-bound IRK contained 56598 atoms. Following energy minimization and equilibration in the NVT (500 ps) and NPT (10 ns) ensembles, three independent 1 *µ*s simulations were conducted. Finally, the conformational analysis and calculation were performed on a total of 6 *µ*s of simulation data from the apo-IRK and the inhibitor-bound IRK. We provide additional details on MD simulations and analyses in the supporting information.

### 2.4 Conformational Analyses

We visualized and analyzed MD simulation trajectories using the GROMACS modules and VMD. The conformational dynamics of the IRK structures was evaluated using multiple metrics. We calculated the root mean squared deviation (RMSD) of the protein backbone atoms taking the initial structure as a reference and the root mean squared fluctuation (RMSF) was evaluated for the C_*α*_ atom of each residue from its mean position. The RMSD was calculated for the heavy atoms of the inhibitor, taking the initial docked conformation as a reference. The interactions between the IRK and the allosteric inhibitor were evaluated using the Protein-Ligand Interaction Profiler (PLIP) web tool [60]. The interactions were visualized, and the images were rendered using PyMol [61]. We further performed, principal component analysis (PCA) to examine conformational transitions and principal motions in the apo-IRK and inhibitor-bound IRK structures. A covariance matrix was generated from eigenvectors representing atomic fluctuations in the protein backbone atoms, which was diagonalized to obtain eigenvectors and eigenvalues. The principal components (PCs) with the highest eigenvalues represent the most significant structural motions [62]. The essential subspace of conformations was defined by projecting onto the first two PCs, PC1 and PC2. We also analyzed the lowest energy conformational ensembles using the Free Energy Surface (FES) projected along the principal components.

## 3 RESULTS

### 3.1 Conformational dynamics revealed a type III allosteric pocket in IRK

We performed MD simulations of the apo IRK structure to understand the structural variations specifically around the potential allosteric pocket region (Figure 1B). We conducted principal component analysis on conformations generated by MD trajectories to extract the principal conformational modes from each trajectory. We assessed the dominant conformational ensembles of the apo IRK structure by projecting the free energy surface (FES) along the first two principal components (PC1 and PC2) (Figure 2). Further, we analyzed protein conformations corresponding to the freeenergy minima and evaluated conformational changes occurring in the IRK structure relative to its initial configuration (X-ray structure of apo IRK inactive conformation; PDB ID 1IRK). We observed variations in key structural elements of a kinase: the *α*C-helix, GC-loop, A-loop, and C-loop (Figure 3A). In MD-derived conformations, we observed an oblique inward displacement of the *α*C-helix relative to its initial conformation. The displacement of the *α*C-helix was correlated with the conformation of the GC-loop (Figure 3A). As the *α*C-helix shifted in a more oblique and inward direction, the GC-loop exhibited an outward movement. Furthermore, the A-loop shows a change in its conformation by forming a short intermediate helical structure (Figure 3A). This helix in the A-loop was formed by the residues, M1153, T1154, R1155, D1156, and I1157, adjacent to the DFG motif (D1150, F1151, and G1152). Moreover, we have previously studied and reported that a helical intermediate state of the A-loop facilitates the “DFG flip” in the IRK [63]. We observed that the DFG motif was stable in the “DFG-out” conformation in all MD trajectories.

**FIGURE 2.**
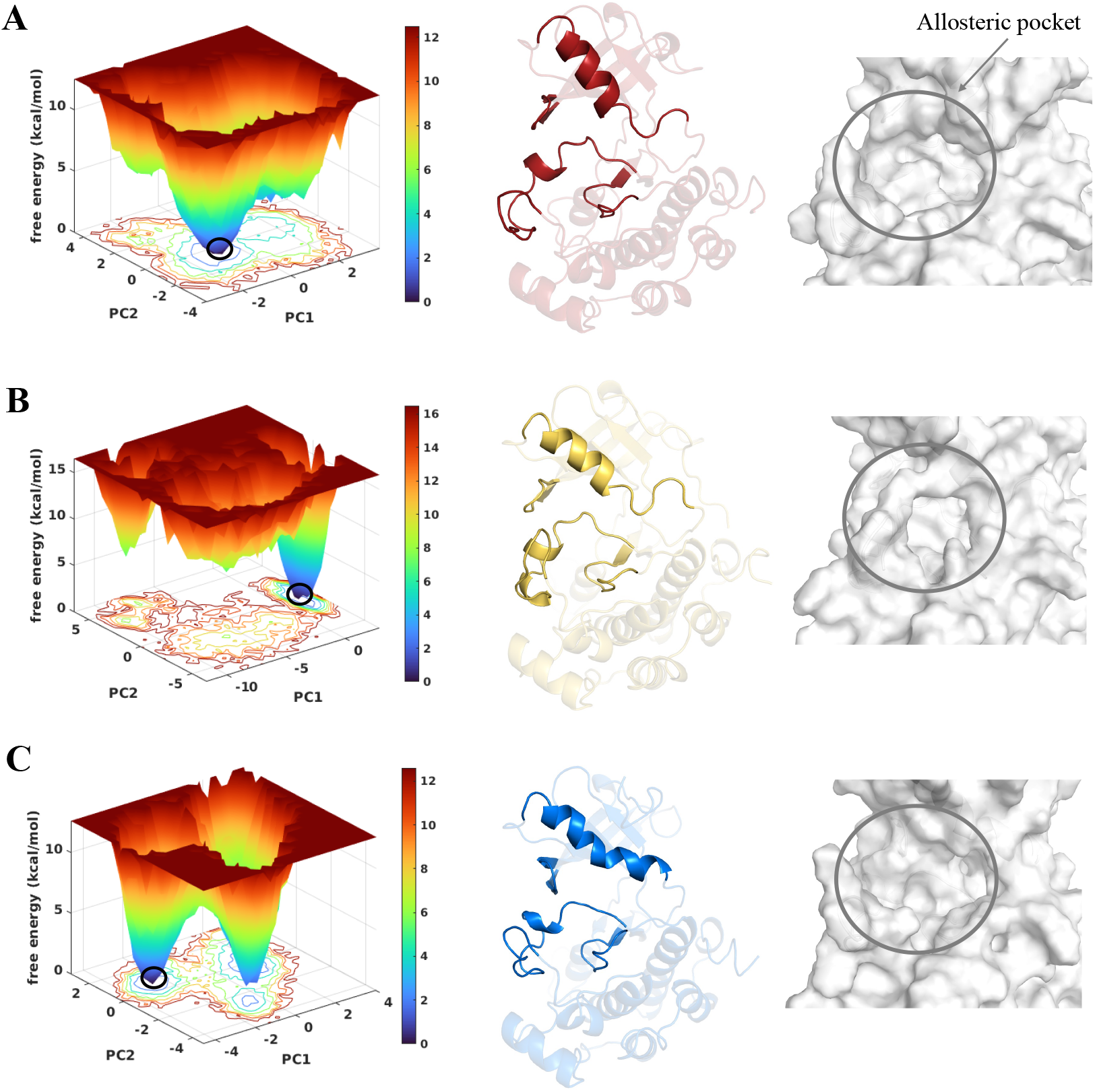
Free energy surface (FES) for apo IRK projected along two principal components (PC1 and PC2) for each of the three independent MD simulation trajectories. (A-C) (left) FES projected along PC1 and PC2; (middle) apo IRK conformation corresponding to the minima circled in the FES plot; and (right) A surface view of the allosteric pocket region.

**FIGURE 3.**
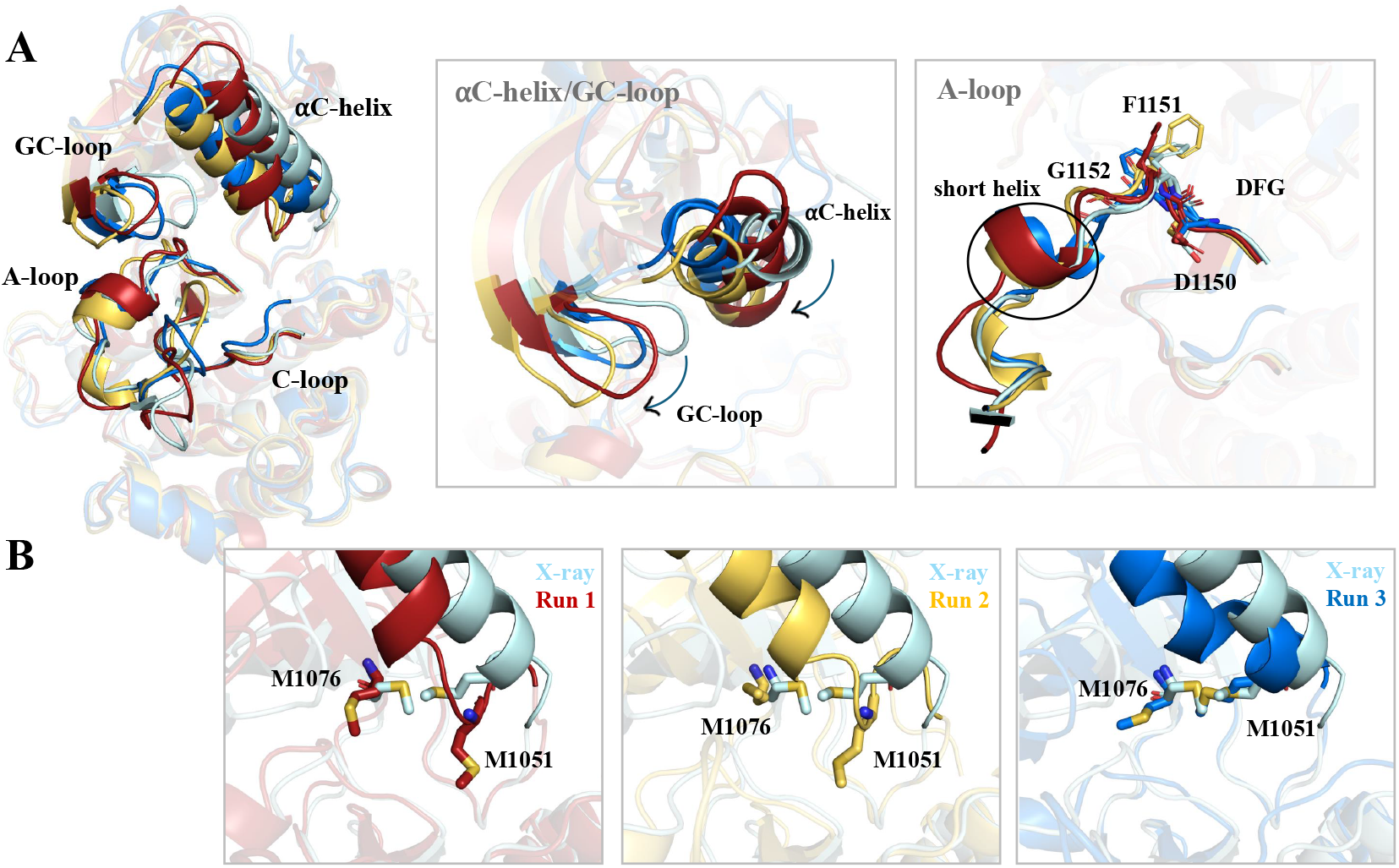
Structural superposition of the MD-derived apo IRK conformations obtained from different free energy surfaces (cf. Figure 2). (A) (left) Cartoon representation of apo IRK conformations: X-ray PDB ID 1IRK (Cyan); Run 1 (Red); Run 2 (Yellow); Run 3 (Blue); (middle) A comparison of *α*C-helix and GC loop conformations; (right) Comparison of the activation loop (A-loop) and the DFG motif residues. (B) A comparison of side chain conformations of residues M1051 and M1076.

In our previous work [47, 64], we have reported conformational dynamics associated with the allosteric pocket residues in IGF1RK responsible for the selectivity of a type III allosteric inhibitor. We described that in contrast with the IG1RK residues M1054 and M1079, the corresponding residues M1051 and M1076 in IRK have strikingly different conformational dynamics [47]. Furthermore, in the current study, we observed that the IRK *α*C-helix partially uncoils from its C-terminal region facilitating a conformational change in the residue M1051 (Figure 3B). These structural variations affect the shape and size of the allosteric pocket and may have a crucial role in recognition of the type III allosteric inhibitors of IRK. These observations indicate that in IRK the *α*C-helix has to uncoil from the C-terminal region for the methionine side chain (M1051) to adopt a conformation similar to M1054 in the IGF1RK allosteric pocket [47, 46]. Consequently, our findings suggest that two homologous proteins with conserved allosteric pocket residues may differ in conformations adopted by the pocket residues, which is crucial for achieving inhibitor selectivity in allosteric inhibition of isoforms and homologs.

The structural deviations in the *α*C-helix, A-loop, GC loop, and C-loop contributed to the changes in the overall size and shape of the allosteric pocket region in IRK. Further, we calculated the structural metrics, RMSD and RMSF of the apo-IRK structure to evaulate the deviations corresponding to these regions (Figure S1). The average RMSD of the backbone atoms of *α*C-helix was about 3Å. We observed this conformational change in *α*C-helix in two trajectories within the first 200 ns, whereas conformational change in the third trajectory was evident after 600 ns (Figure S1B). The A-loop in all three trajectories showed higher deviations of the backbone atoms with an RMSD value of about 4.5 Å(Figure S1B). The GC loop, however, deviates in coordination with the movement of the *α*C-helix. Higher deviations were observed toward the end of the trajectory (Figure S1B). We further evaluated the RMSF for the C_*α*_ atoms of the residues, to characterize each region with a higher flexibility (Figure S1B). There was an evident peak showing higher fluctuation corresponding to the A-loop (residues 1149-1170). The RMSF for the *α*C-helix (1037 to 1053) residues at the N-terminus of the helix were slightly higher (1.5 Å) than the residues at the C-terminus (1 Å). We also observed uncoiling of the *α*C-helix in the C-terminal region in two MD trajectories. The uncoiling was coupled with the conformational change in the sidechain of the residue M1051 in the *α*C-helix (Figure 3B). Thus, our results support that the type III allosteric pocket aka “back pocket” in IRK is accompanied by structural variations influencing the overall conformation of the allosteric region. Hence, to further validate and characterize this type III allosteric site in IRK, we analyzed the features of this pocket throughout our MD trajectories.

### 3.2 Pocket analysis uncovered a stable non-transient allosteric site

We utilized a geometric approach based on Voronoi tessellation to identify potential cavities in apo IRK conformational ensembles based on MD trajectories. We aimed to assess and examine the variations associated with the “back pocket” region. The frequency and density grid maps obtained using the MDpocket tool were analyzed to characterize and understand the shape and physicochemical properties of the allosteric pocket region. These maps were examined and visualized using VMD. For each MD trajectory, both the frequency and density grid map were generated and analyzed independently. In Figure 4A, we show the frequency grid maps for each trajectory, representing the pockets consistently observed during trajectories. The uniquely colored mesh regions for each trajectory correspond to the cavities most frequently open and accessible during the trajectory. The frequency grid map shown in Figure 4A has a threshold of isovalue 0.5; hence, focusing on grid points that are consistently occupied by *α* -spheres across the trajectory, thereby highlighting more reliable and recurrent binding sites. Here, the encircled area in the frequency grid maps represents the potential allosteric pocket region in IRK, suggesting a higher frequency of its occurrence. Furthermore, the density grid maps for each trajectory are shown in Figure S2. We observed multiple cavities/pockets in the density grid map of the IRK structure, identified across the simulation trajectories. The density grip maps detects the transient channels as well as more conserved cavities. We observed that the region between the *α*C-helix and the A-loop has a higher atomic density, representing the occurrence of potential binding sites or pockets. This region in IRK is behind the ATP binding pocket also known as the “back pocket” (type III pocket). Furthermore, this region was identified in the density grid map even at higher isovalues, suggesting that the allosteric pocket is more conserved and dense (Figure S2). Therefore, the type III allosteric pocket region in IRK has a higher frequency of occurrence, Given that pocket druggability can be influenced by conformational changes within the binding pocket, it is essential to contextualize these dynamic alterations and correlate them with relevant physical or chemical descriptors. We evaluated several descriptors for the allosteric pocket region by analyzing each trajectory and averaging the descriptors across all simulations. As the physicochemical properties of the pocket are largely determined by residues surrounding the pocket, we first identified these residues involved in the formation of the allosteric pocket. Key binding site residues were defined as those consistently present across all MD frames, based on their proximity to the grid-defined pocket (Figure 4B). Since proteins are dynamic entities, we observed notable changes in the pocket size and shape, signifying changes in the pocket volume and density. This observation was corroborated by density and frequency maps, indicating that this pocket is among the largest and most frequently observed among all protein cavities throughout each trajectory. The average volume of the type III allosteric pocket in IRK was 751.4 Å^3^ (Figure 4C). Overall, the averaged (over 3 simulations) volume of the cavity remains consistent. The density plot shows an increase in the density toward the end of the simulation trajectory, reflecting a higher number of atoms present in the pocket (Figure 4C). The mean local hydrophobic density (MLHD) is an important descriptor, which indicates the presence of hydrophobic amino acid residues and the associated compactness within the pocket. It reflects local densities of hydrophobic *α* sphere clusters within the cavity (Figure 4C). A higher MHLD value indicates that the pocket will favor interactions with small hydrophobic drug-like compounds by increasing the possibility of stronger protein-ligand interactions through a hydrophobic effect. The average MLHD for the allosteric pocket in IRK was 47.8. MLHD is an essential metric while comparing and identifying drug-binding pockets as it is correlated with the druggability of the binding pocket. Furthermore, the average solvent-accessible surface area (SASA) for the pocket was 453.5 Å^2^ (Figure 4C). Additionally, we determined the electrostatic potential map of the IRK structure to identify and compare the charge distribution of the orthosteric (ATP binding pocket) and the type III allosteric binding pocket. The electrostatic potential map of IRK revealed a higher negative potential at the ATP binding pocket (Figure S3). Similarly, for the type III allosteric pocket in IRK, we observed a higher negative electrostatic potential at the core of the pocket, with lower electrostatic potential in the surrounding region (Figure 4D). Thus, an allosteric modulator with positively charged or polar functional groups could be developed to target the negatively charged core of the type III allosteric pocket of IRK.

**FIGURE 4.**
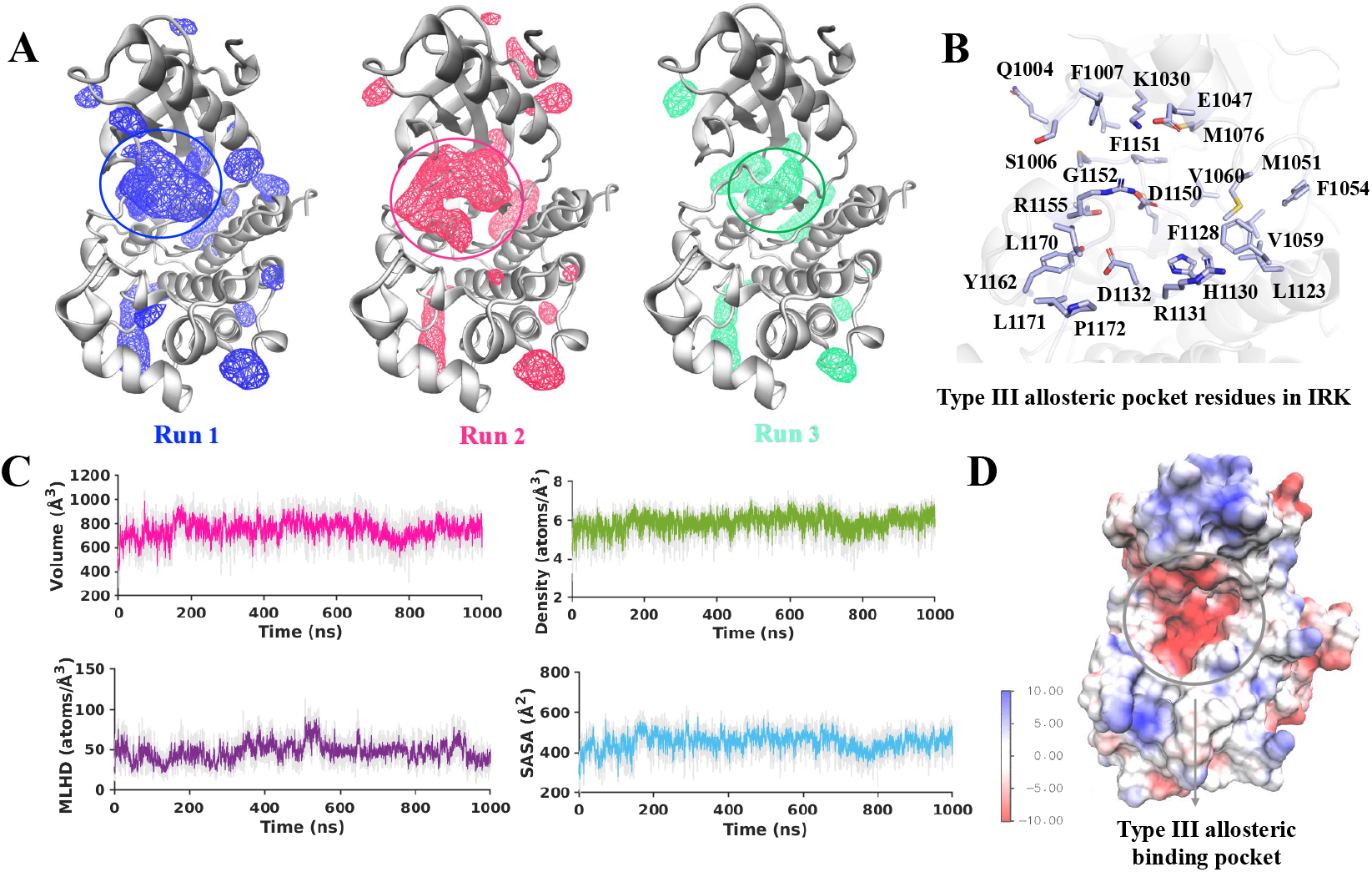
(A) Frequency grid maps obtained for each of the three independent MD simulation trajectories. The grid maps shown have a threshold of isovalue 0.5, with the encircled grid highlighting the allosteric pocket region. (B) IRK residues involved in the formation of the type III allosteric pocket. (C) Physiochemical properties of the selected allosteric pocket derived from the frequency grid maps. Each property is averaged over three trajectories: Volume, Density, Mean Local Hydrophobic Density (MLHD), and Solvent Accessible Surface Area (SASA). (D) Electrostatic potential energy surface map of the IRK domain. suggesting a non-transient stable binding pocket (encircled in Figure 4A).

### 3.3 An allosteric inhibitor stablizes the inactive conformation of *α*C-helix

We modeled the interaction of an experimentally known IRK inhibitor at the identified allosteric binding pocket in IRK (Figure 5). From a series of indole-butyl-amines derivatives, the most potent derivative against IRK was selected for docking studies [46]. In the initially docked conformation, one of the indole rings (indole ring R1) of the inhibitor is positioned at the shallow hydrophobic region below the *α*C-helix (Figure S4B). The indole ring R2 interacts with the A-loop and stacks parallel to the C-loop. The inhibitor binding to the type III allosteric pocket or the “back pocket” in IRK will likely impede the conformational change associated with the inactive and active conformation of the IRK domain.

**FIGURE 5.**
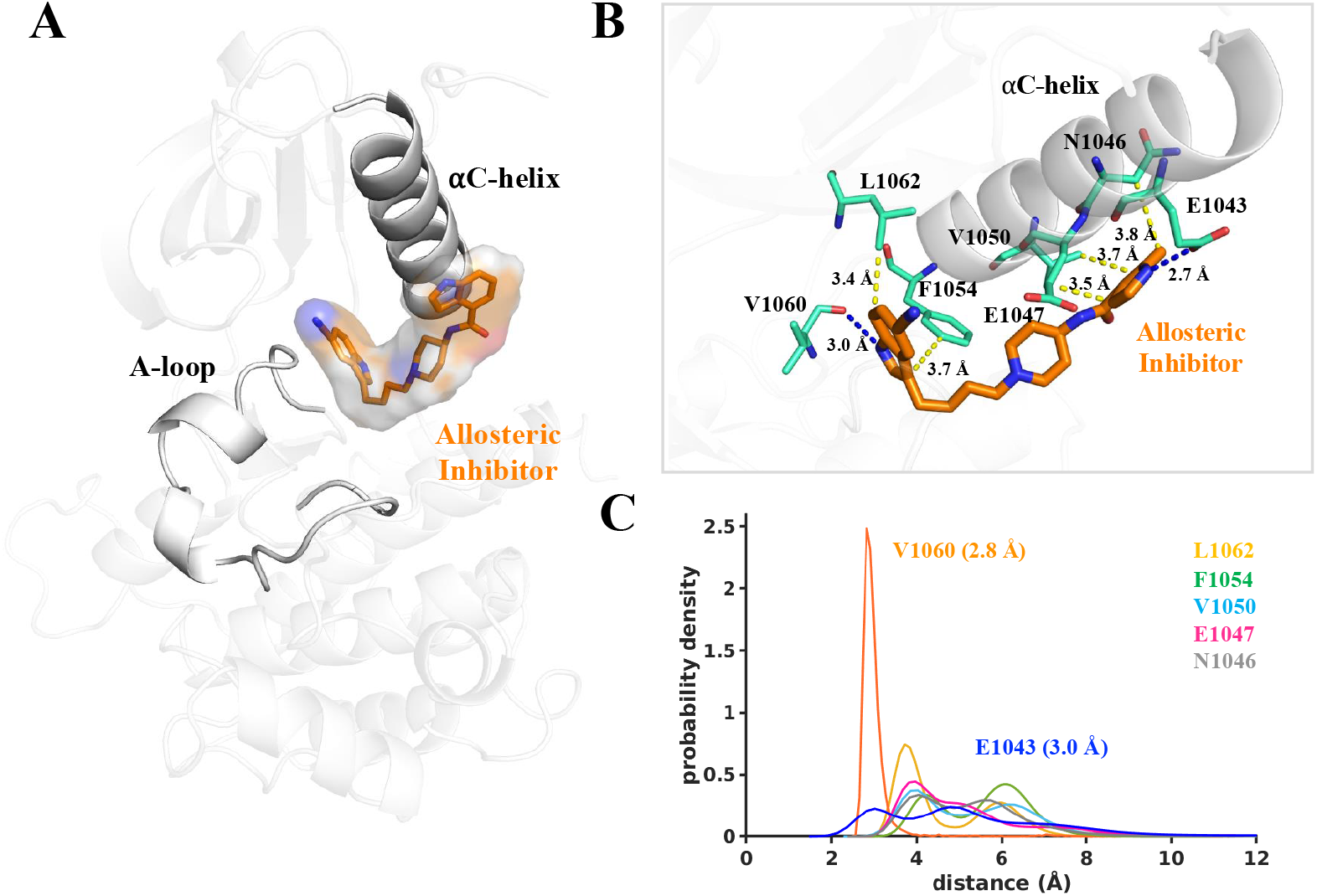
(A) The MD derived conformation of IRK bound to the allosteric inhibitor at the type III pocket. (B) Binding conformation and interactions of the allosteric inhibitor with the IRK residues. The hydrogen bonds are shown with blue dashed lines and hydrophobic interactions are shown in yellow. (C) The distribution of distance for each interaction between the IRK residues and the allosteric inhibitor. The distribution data are based on three independent MD runs. Each curve is represented by a unique color for its corresponding residue. The distribution peaks for hydrogen bonds are observed at 2.8 Å for residue V1060 and 3.0 Å for residue E1043. The peaks for other residues are as follows: L1062 (3.7 Å and 5.9 Å); F1054 (4.2 Å and 6.1 Å); V1050 (3.85 Å); E1047 (3.8 Å); N1046 (4.0 Å).

To further assess the stability of the inhibitor within the binding pocket, we performed three independent MD simulation runs to evaluate the flexibility of the complex. Based on 3 *µ*s of simulation data, we then identified the predominant conformation of the inhibitor within the type III allosteric pocket of IRK by analyzing over 3 *µ*s of MD simulation data. We assessed the RMSD of the non-hydrogen atoms of the inhibitor. The RMSD data indicate higher deviations of the inhibitor atoms in comparison to their positions in the docked conformation (initial conformation), with the deviation reaching a plateau after 400 ns (Figure S5). This observation suggests that the inhibitor initially deviates from its docked conformation and subsequently adopts a stable conformation after 400 ns. All three trajectories exhibited a similar trend and showed comparable overall binding conformations of the inhibitor within the allosteric pocket (Figure S5). The dominant conformation of the inhibitor in the IRK allosteric pocket features its indole ring stacked parallel to the *α*C-helix (Figure 5A). The inhibitor undergoes an approximate 3 Å deviation from its initial conformation to achieve this stable orientation. Notably, the indole-4-carboxamide substituent interacts with the *α*C-helix residues E1043, N1046, E1047, and V1050 (Figure 5B). A detailed interaction map of the inhibitor within the binding pocket is shown in Figure 5B. The hydrogen bonding interaction with the residue V1060 remains stable throughout the simulation trajectory, displaying a single distribution peak at 2.8 Å (Figure 5C). Furthermore, the probability density distribution data reveals that the hydrogen bond between E1043 and the nitrogen donor atom of the inhibitor is observed at distances of 3 Å and 4.7 Å. The hydrophobic interactions with residues N1046, V1050, E1047, F1054, and L1062 also contribute to stabilizing the inhibitor in its conformation.

Furthermore, we performed a comprehensive analysis of the residues forming the type III allosteric pocket in IRK. To estimate the flexibilities of the residues in the binding pocket, we calculated the RMSF for each residue in both the inhibitor-bound and unbound forms (Figure 6A). The RMSF values were derived by averaging over three simulation trajectories for each state. The residues S1004, Q1006, and F1007 from the GC-loop exhibited higher fluctuations (> 1.5 Å) in the inhibitor-bound form compared to the unbound state. In contrast, residues E1043, E1047, M1051, and F1054 from the *α*C-helix region displayed significantly lower flexibility when the inhibitor was bound, indicating that interactions between these residues and the inhibitor might contribute to their reduced fluctuations. The A-loop residues, including the DFG motif, demonstrated slightly increased flexibility in the inhibitor-bound form. However, the presence of the inhibitor at the allosteric binding pocket allows the residues to adopt conformations that may facilitate potential interactions. Furthermore, the chemical characteristics of the binding pocket residues governs the conformation and interactions of the inhibitor at the allosteric pocket. We categorized residues in the allosteric binding pocket into polar, non-polar (aromatic and aliphatic), and charged groups (Figure 6B). Hence, we generated a chemical characteristic map of the type III allosteric pocket of IRK (Figure 6C).

**FIGURE 6.**
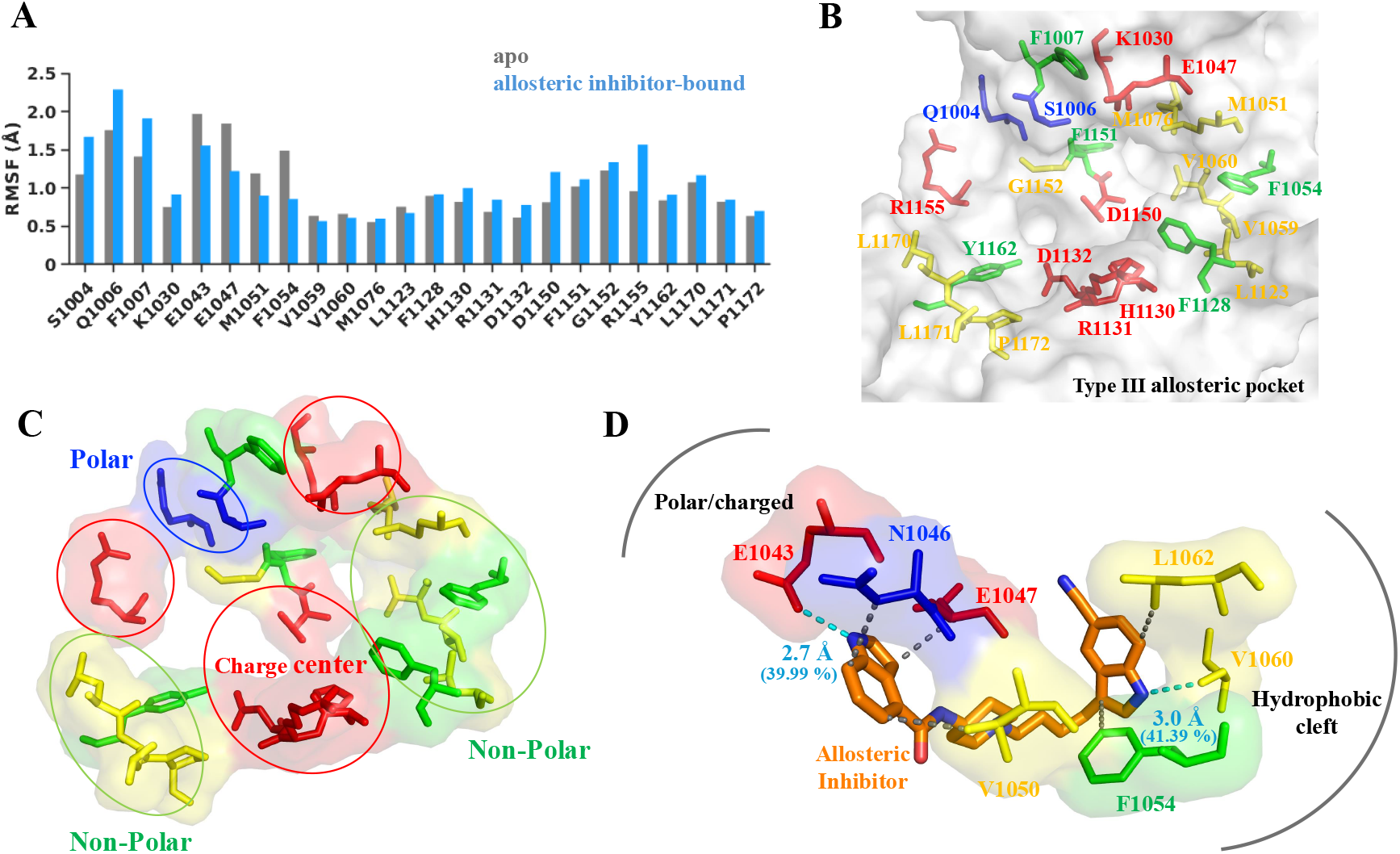
(A) The root mean squared fluctuations (RMSF) of the C*α* atoms of allosteric pocket residues in both apo and inhibitor-bound IRK states. The RMSF values are averaged over three MD simulation trajectories. (B) The type III allosteric pocket residues in the IRK. (C) A chemical characteristic map of the type III allosteric pocket of IRK. (D) Interactions between the inhibitor and the allosteric pocket residues. The hydrogen bonds are labeled with their distances and their occupancies. The allosteric pocket residues in panels (B), (C), and (D) are classified into three categories: polar, non-polar (aromatic and aliphatic), and charged. Each category is color-coded as follows: polar: blue; charged: red; non-polar (aliphatic): yellow; non-polar (aromatic): green.

The type III allosteric pocket features a hydrophobic core composed of both aromatic and aliphatic non-polar residues, forming a cleft-like region (Figure 6D). The DFG and HRD motifs, along with the catalytic residues K1030 and E1047, create a charge center, while polar residues are positioned towards the outer surface. Additionally, another non-polar region is formed by the terminal A-loop residues. The inhibitor in the allosteric pocket orients toward the *α*C-helix; this conformation was observed stable in all MD trajectories. The cyano-indole ring of the inhibitor interacts with the hydrophobic cleft, stabilized by a persistent hydrogen bond with V1060 (Figure 6D). Meanwhile, the indole-4-carboxamide substituent is oriented toward the polar and charged residues of the *α*C-helix, forming a hydrogen bond with E1043 (Figure 6D). Thus, the inhibitor acquires a conformation in the allosteric pocket that stablizes the inactive conformation of the *α*C-helix.

### 3.5 Type III allosteric pocket in IRK has *α*C-helix “out”/DFG “out” conformation

To further comprehend the structural aspects of the key kinase elements, we compared the MD derived conformations of the IRK structures (apo and inhibitor-bound) to the X-ray crystal structures of the inactive (PDB ID 1IRK) and active (PDB ID 1IR3) IRK (Figure 7). We performed a structural alignment based on the C_*α*_ atoms of the following structures: IRK inactive conformation (PDB ID: 1IRK), IRK active conformation bound to ATP (PDB ID: 1IR3), MD-derived conformation of the IRK apo form (inactive), and MD-derived conformation of the allosteric inhibitor-bound form. The coordinates from the PDB ID 1IRK were used as a point of reference for structural superposition. We investigated the key structural elements surrounding the type III allosteric binding pocket region in IRK. We report that the type III allosteric site in IRK is sandwiched between the N-terminal and the C-terminal lobes, beneath the *α*C-helix and the beta-sheets of the N-terminal domain. Our results indicate that this pocket is present in the inactive IRK and is significantly influenced by the conformation of the *α*C-helix and the A-loop.

**FIGURE 7.**
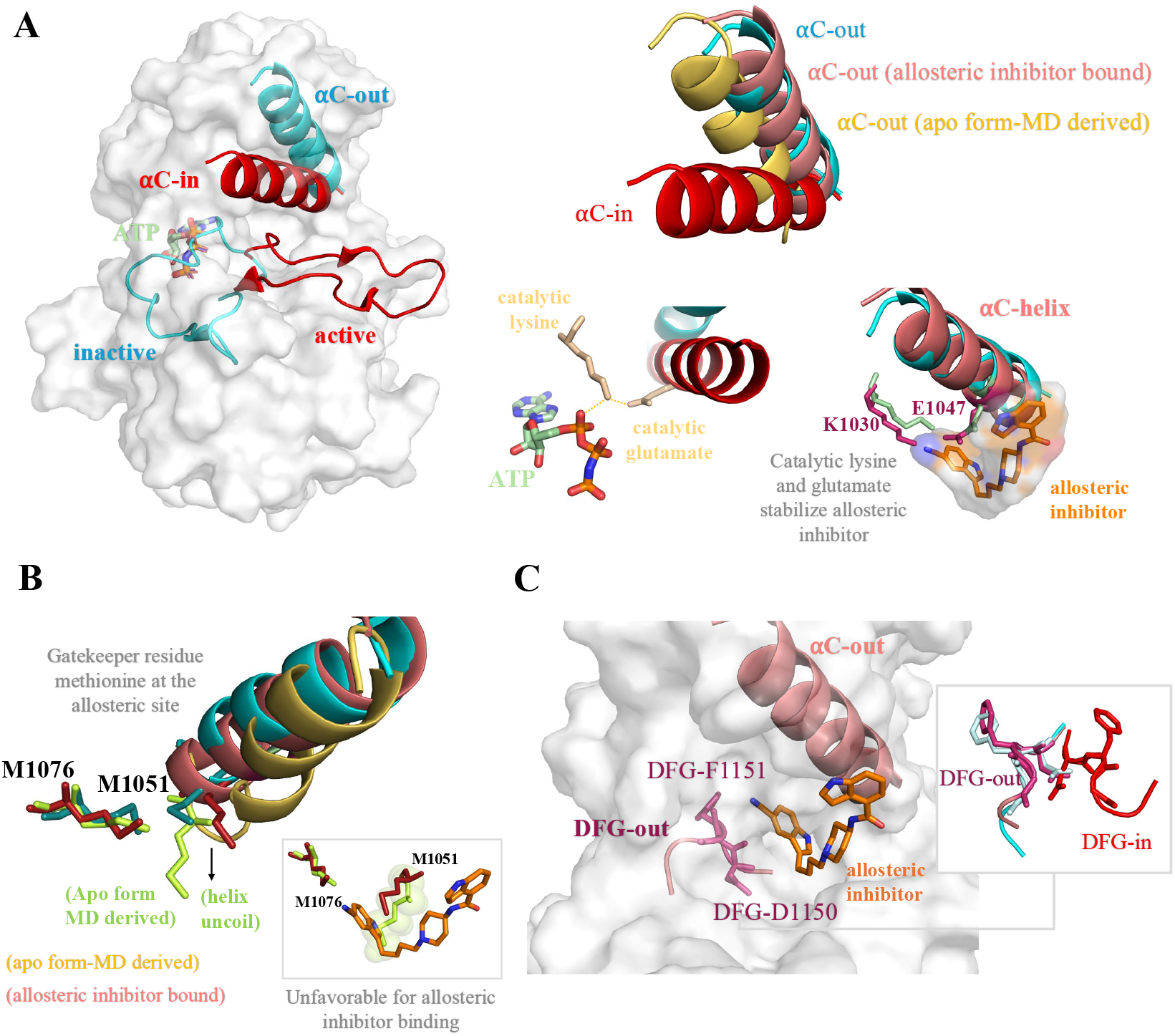
Conformations of the key structural elements at the type III allosteric pocket in IRK. (A) (left) A structural comparison of the *α*C-helix and the activation loop (A-loop) conformations in the inactive IRK (PDB ID 1IRK) and active IRK (PDB ID 3IRK). (right-upper panel) Conformations of the *α*C-helix in multiple IRK states.(right-lower left panel) Interaction between the catalytic lysine and glutamate with ATP bound in the active IRK state (PDB ID: 3IRK). (right-lower right panel) Conformation of the catalytic lysine (K1030) and glutamate (E1047) in the presence of an allosteric inhibitor bound to IRK. (B) A comparison of the sidechain of residues M1051 and M1076 from different IRK conformations. (C) The conformation of DFG motif in the allosteric inhibitor-bound IRK. The right panel displays the DFG-in (active IRK) and DFG-out (inactive IRK) conformations for reference

Protein kinases are known to be regulated through conformational shifts in two structural elements, the *α*C-helix and the A-loop (Figure 7A). In the active IRK, the A-loop adopts an open conformation, while the *α*C-helix is oriented inward, “*α*C-in”. In contrast, the inactive IRK has its A-loop folded inwards and the *α*C-helix is in “*α*C-out” conformation (Figure 7A). Furthermore, the structural comparison suggests that the *α*C-helix conformation observed in the MD-derived conformation of the IRK apo form has an inward oblique shift (Figure 7A) (also discussed in the previous section (Figure 3). In contrast, the allosteric inhibitor bound structure exhibits a conformation of the *α*C-helix which is comparable to that observed in the inactive state (Figure 7A). The *α*C-helix is orientated to facilitate substrate binding in the active conformation, and this state is stabilized by a crucial contact between the catalytically important lysine and glutamate residues (Figure 7A). However, inhibitor binding to the type III allosteric pocket in the IRK can promote a conformation of the *α*C-helix that disrupts this interaction, leading to the inactive state (Figure 7A). Notably, the residues E1047 and K1030 participate in interactions with the inhibitor that help stabilize this conformation. Consequently, the disruption of the salt bridge between the catalytic glutamate and lysine serves as a strong indicator of protein kinase inactivity, highlighting a mechanism of allosteric regulation. These observations suggest that IRK can undergo allosteric inhibition, marked by changes in the *α*C-helix conformation and interactions between critical amino acids.

Furthermore, two methionine residues, M1051 and M1076, play essential roles in the overall conformation of the allosteric pocket and inhibitor binding. The conformation of the residue M1051 also influences the overall conformation as well as the secondary structure of the *α*C-helix. Moreover, M1051 act as a gatekeeper residue, as its sidechain conformation impacts the binding of the allosteric inhibitor within the hydrophobic core of the allosteric pocket. In apo IRK, the sidechain of M1051 is oriented such that it creates steric hindrance, thereby obstructing the binding of the inhibitor, particularly the cyano-indole ring (Figure 7B). Conversely, when the inhibitor is bound, the M1051 sidechain adopts a conformation that stabilizes the inhibitor in the allosteric pocket (Figure 7B). Hence, these results highlight the significance of residue conformations in modulating the type III allosteric inhibitor binding in IRK.

The conformation of the DFG motif is another key indicator of kinase activity. In the DFG motif-”in” conformation, the D1150 residue (in the context of IRK) is positioned to facilitate a catalytically competent active conformation. In contrast, in the “out” conformation of the DFG motif, the D1150 residue points away from the ATP binding site, while the phenylalanine residue F1151 occupies the ATP pocket, resulting in a catalytically inactive conformation (Figure 7C). When the inhibitor occupies the type III allosteric binding pocket in IRK, the DFG motif is restricted in the “out” conformation. The D1150 residue intermittently participates in hydrogen bonding interaction with the piperidine ring of the allosteric inhibitor. Furthermore, the orientation of the cyano-indole ring of the inhibitor induces steric hindrance which prevents the DFG-flip necessary for adopting an “in” conformation, thereby restricting the DFG motif in its “out” conformation. Consequently, the type III allosteric pocket in IRK is characterized by a “DFG-out” conformation.

### 4 DISCUSSION

Exploiting conformational variations in protein kinase structures has proven to be extremely valuable for developing kinase inhibitors [65, 66, 67]. Moreoever, computational approaches that capture protein dynamics provide an unprecedented opportunity for discovering novel allosteric sites and allosteric modulators with a desired selectivity profile [29]. In our previous study [47], we employed a suite of computational methods to provide a comprehensive analysis of allosteric inhibitor binding in IGF1RK and IRK. In a subsequent study [64], we established structural models of experimentally known allosteric inhibitors of the insulin receptor family of kinases. Considering that the type III allosteric pocket is present in the IRK, we have characterized the structural features specifically associated with this allosteric site.

Our results from the conformational dynamics analysis revealed variations in key structural elements surrounding the allosteric pocket region, particularly in the *α*C-helix, the A-loop, and the GC loop. Additionally, the application of a geometric method-based approach allowed us to predict and identify cavities corresponding to the allosteric pocket region in the IRK structure, providing insights into the spatial characteristics of this critical site. The density and frequency maps indicated that the region between the *α*C-helix and the A-loop has high atomic density and is highly conserved, suggesting a stable and accessible “back pocket”. Our analysis of the chemical characteristics of this allosteric pocket involved calculating descriptors of the selected pocket region, which helped validate and identify its physicochemical features. We identified residues that consistently form the allosteric pocket, providing insight into its structural composition. The observed increase in volume and density of the pocket during the simulation indicates dynamic conformational changes, which may enhance its potential for inhibitor binding. Additionally, the hydrophobic density curve and the electrostatic potential map further suggested a favorable environment for potential inhibitor interactions. Our results reveal that the allosteric pocket is characterized by a hydrophobic groove composed of both aliphatic and aromatic non-polar residues, along with a charge center that facilitates electrostatic and ionic interactions. The high negative electrostatic potential at the core, coupled with the lower potential surrounding the pocket, suggests that a type III allosteric modulator targeting this site would likely require a hydrophobic group to interact with the non-polar residues, along with a positively charged group to engage with the electrostatically favorable core. This detailed chemical characterization of the type III pocket in IRK provides essential insight for guiding the rational design of structure-based allosteric modulators, advancing the development of more specific and effective therapeutic agents.

Building on our findings, we investigated the binding of an experimentally known IRK inhibitor (N-1-[4-(5-cyano-1H-indol-3-yl)butyl]piperidin-4-yl-1H-indole-4-carboxamide) to the allosteric pocket. This inhibitor represents the most potent compound against IRK with an IC50 value of 1.3 *µ*M, as reported in a series of allosteric modulators targeting the insulin family of kinases [46]. Consistent with our results, which proposed that an allosteric modulator targeting this site would likely require both a hydrophobic group and a positively charged group, the structure of this inhibitor aligns with our hypothesis. Specifically, the inhibitor features indole rings with a long acyl chain as the hydrophobic moiety, while groups such as the carbonyl oxygen and amino group facilitate electrostatic interactions at the allosteric pocket. This supports the notion that the most potent inhibitor in this series aligns with the structural features we proposed for optimizing the binding to the type III allosteric pocket in IRK. Furthermore, our results suggest allosteric inhibitor binds stably, and its interactions with the *α*C-helix demonstrate its ability to stabilize the inactive conformation of IRK. Notably, the stable hydrogen bonding interactions with the residues E1043 and V1060 restrict the movement of the *α*C-helix, maintaining it in the “out” conformation. Additionally, the RMSF analysis of pocket residues indicate reduced flexibility of the *α*C-helix in the inhibitor-bound form, suggesting that inhibitor binding stabilizes this region. In contrast, residues in the A-loop exhibit increased flexibility, which could indicate potential dynamic interactions with the inhibitor, further supporting the idea of a flexible binding site.

Moreover, the structural comparison between the apo and inhibitor-bound IRK conformations allowed us to characterize the key structural elements, specifically around the allosteric binding pocket. Notably, we observed an oblique inward displacement of the *α*C-helix correlated with the outward movement of the GC loop, emphasizing their interdependence in the apo IRK conformations. Conversely, in the inhibitor-bound form, the interactions stabilize the *α*C-helix, adopting an “out” conformation. Our findings imply that the allosteric inhibitor binding at the type III pocket promotes an “out” conformation of *α*C-helix, thereby preventing ATP interaction with catalytic residues K1030 and E1047 in IRK, which is essential for catalytic activity of the kinase. The allosteric modulation of various kinases is associated with the *α*C-helix displacement at the allosteric pocket [45]. The displaced *α*C-helix provides space for ligand binding, and this *α*C-helix “out” conformation is stable regardless of the presence of ATP-site binders. Thus, our study reports that the allosteric inhibitor binding in the type III pocket of IRK will promote the “out” conformation of the *α*C-helix. Our results show that in the apo state of IRK, the sidechain of M1051 adopts a conformation that induces the uncoiling of the *α*C-helix from its C-terminal end and consequently leading to an unfavorable conformation of the allosteric pocket for inhibitor binding. Conversely, in the inhibitor-bound conformation, the *α*C-helix remains intact, and M1051 adopts a conformation that stabilizes the inhibitor within the pocket. Our study thus suggests that M1051 functions as a gatekeeper residue, crucial for maintaining the structural integrity of the *α*C-helix and regulating the binding of the allosteric inhibitor. Furthermore, our findings show that the A-loop forms a short helical intermediate adjacent to the DFG motif. The DFG motif, highly conserved in kinases, plays a critical role in kinase activity, and previous studies have demonstrated that type III allosteric inhibitors preferentially stabilize the “DFG-out” conformation [68, 69, 42]. Consistent with this, we observed that inhibitor binding to the type III allosteric pocket in IRK stabilizes the “DFG-out” conformation. Overall, our findings provide an understanding of the structural and molecular details of the type III allosteric pocket of IRK for further development of allosteric modulators with higher specificity and selectivity for the insulin receptor family of kinases.

### 5 CONCLUSION

Overall, this study unveils the conformational dynamics and structural features of the type III allosteric pocket in IRK, offering insights for the development of selective inhibitors. Our results suggest that the type III pocket in IRK is an accessible back pocket beneath the *α*C-helix and *β* -sheets of the N-terminal lobe, with a favorable environment for inhibitor binding. The pocket features a hydrophobic cleft, with a core exhibiting high electrostatic potential and charged residues, conducive to interaction with potential modulators. Our data confirm that the *α*C-helix is stabilized in its “out” conformation by persistent interactions with the allosteric modulator, thereby maintaining the inactive conformation of IRK. Furthermore, the DFG motif in the A-loop adopts a “DFG-out” conformation in both apo and inhibitor-bound IRK. Additionally, our study identifies M1051 as a critical gatekeeper residue essential for the structural integrity of the *α*C-helix and the regulation of allosteric inhibitor binding. Together, these findings provide valuable insights into the conformational dynamics of key structural motifs in the IRK allosteric binding site, aiding the design of selective allosteric modulators for the targeted inhibition of receptor tyrosine kinases.

## acknowledgements

We acknowledge the financial support provided by the National Institutes of Health (NIH) through Grant R35GM138217. The content is solely the responsibility of the authors and does not necessarily represent the official views of the NIH. We are grateful for computational support through Premise, a central shared HPC cluster at UNH supported by the Research Computing Center.

## conflict of interest

The authors declare no conflicts of interest.

## supporting Information

Additional supporting information can be found online in the Supporting Information section at the end of this article.

## Notes

### Competing Interest Statement

The authors have declared no competing interest.

## references

[1] Hunter T. Protein kinases and phosphatases: The Yin and Yang of protein phosphorylation and signaling. Cell 1995;80(2):225–236.

[2] Ubersax JA, Ferrell Jr JE. Mechanisms of specificity in protein phosphorylation. Nat Rev Mol Cell Biol 2007;8(7):530– 541.

[3] Cheng HC, Qi RZ, Paudel H, Zhu HJ. Regulation and Function of Protein Kinases and Phosphatases. Enzyme Res 2011;2011(1):794089.

[4] Manning G, Whyte DB, Martinez R, Hunter T, Sudarsanam S. The Protein Kinase Complement of the Human Genome. Science 2002;298(5600):1912–1934.

[5] Duong-Ly KC, Peterson JR. The Human Kinome and Kinase Inhibition. Curr Protoc Pharmacol 2013;60(1):2.9.1–2.9.14.

[6] Cohen P. Protein kinases — the major drug targets of the twenty-first century? Nat Rev Drug Discov 2002;1(4):309–315.

[7] Al-Masri C, Trozzi F, Lin SH, Tran O, Sahni N, Patek M, et al. Investigating the conformational landscape of AlphaFold2-predicted protein kinase structures. Bioinform adv 2023 09;3(1):vbad129.

[8] Welsh CL, Conklin AE, Madan LK. Crystal Structures Reveal Hidden Domain Mechanics in Protein Kinase A (PKA). Biology 2023;12(11):1370.

[9] Taylor SS, Wu J, Bruystens JGH, Del Rio JC, Lu TW, Kornev AP, et al. From structure to the dynamic regulation of a molecular switch: A journey over 3 decades. J Biol Chem 2021;296:100746.

[10] Zorn JA, Wang Q, Fujimura E, Barros T, Kuriyan J. Crystal Structure of the FLT3 Kinase Domain Bound to the Inhibitor Quizartinib (AC220). PLOS ONE 2015 04;10(4):1–15.

[11] Hubbard SR, Miller WT. Receptor tyrosine kinases: mechanisms of activation and signaling. Curr Opin Cell Biol 2007;19(2):117–123.

[12] Endres NF, Barros T, Cantor AJ, Kuriyan J. Emerging concepts in the regulation of the EGF receptor and other receptor tyrosine kinases. Trends Biochem Sci 2014;39:437–446.

[13] Cabail MZ, Li S, Lemmon E, Bowen ME, Hubbard SR, Miller WT. The insulin and IGF1 receptor kinase domains are functional dimers in the activated state. Nat Commun 2015;6(1):6406.

[14] Kono SA, Marshall ME, Ware KE, Heasley LE. The fibroblast growth factor receptor signaling pathway as a mediator of intrinsic resistance to EGFR-specific tyrosine kinase inhibitors in non-small cell lung cancer. Drug Resist Updat 2009;12(4):95–102.

[15] Salokas K, Liu X, Öhman T, Chowdhury I, Gawriyski L, Keskitalo S, et al. Physical and functional interactome atlas of human receptor tyrosine kinases. EMBO reports 2022;23(6):e54041.

[16] Arkhipov A, Shan Y, Das R, Endres NF, Eastwood MP, Wemmer DE, et al. Architecture and Membrane Interactions of the EGF Receptor. Cell 2013;152:557–569.

[17] Nagao H, Cai W, Albrechtsen NJW, Steger M, Batista TM, Pan H, et al. Distinct signaling by insulin and IGF-1 receptors and their extra- and intracellular domains. PNAS 2021;118(17):e2019474118.

[18] Hubbard SR, Wei WA Lei Hendrickson. Crystal structure of the tyrosine kinase domain of the human insulin receptor. Nature 1994;372(6508):746–754.

[19] Ofer P, Heidegger I, Eder IE, Schöpf B, Neuwirt H, Geley S, et al. Both IGF1R and INSR knockdown exert antitumorigenic effects in prostate cancer in vitro and in vivo. Mol Endocrinol 2015;29(12):1694–1707.

[20] Belfiore A, Malaguarnera R. The Insulin Receptor: A New Target for Cancer Therapy. Front Endocrinol 2011;2:93.

[21] Buck E, Mulvihill M. Small molecule inhibitors of the IGF-1R/IR axis for the treatment of cancer. Expert Opin Investig Drugs 2011;20(5):605–621.

[22] Carboni JM, Wittman M, Yang Z, Lee F, Greer A, Hurlburt W, et al. BMS-754807, a small molecule inhibitor of insulin-like growth factor-1R/IR. Mol Cancer Ther 2009;8(12):3341–3349.

[23] Evans T, Lindsay CR, Chan E, Tait B, Michael SA, Day S, et al. Phase I dose-escalation study of continuous oral dosing of OSI-906, a dual tyrosine kinase inhibitor of insulin-like growth factor-1 receptor (IGF-1R) and insulin receptor (IR), in patients with advanced solid tumors. J Clin Oncol 2010;28(15_suppl):2531–2531.

[24] Strickler HD, Wylie-Rosett J, Rohan T, Hoover DR, Smoller S, Burk RD, et al. The Relation of Type 2 Diabetes and Cancer. Diabetes Technology & Therapeutics 2001;3(2):263–274.

[25] Coughlin SS, Calle EE, Teras LR, Petrelli J, Thun MJ. Diabetes Mellitus as a Predictor of Cancer Mortality in a Large Cohort of US Adults. Am J Epidemiol 2004;159(12):1160–1167.

[26] Vigneri P, Frasca F, Sciacca L, Frittitta L, Vigneri R. Obesity and cancer. Nutr Metab Cardiovasc Dis 2006;16(1):1–7.

[27] Fair AM, Dai Q, Shu XO, Matthews CE, Yu H, Jin F, et al. Energy balance, insulin resistance biomarkers, and breast cancer risk. Cancer Detection and Prevention 2007;31(3):214–219.

[28] Cohen P, Cross D, Jänne PA. Kinase drug discovery 20 years after imatinib: progress and future directions. Nat Rev Drug Discov 2021;20(7):551–569.

[29] Pan Y, Mader MM. Principles of Kinase Allosteric Inhibition and Pocket Validation. J Med Chem 2022;65(7):5288–5299.

[30] Vijayan RSK, He P, Modi V, Duong-Ly KC, Ma H, Peterson JR, et al. Conformational Analysis of the DFG-Out Kinase Motif and Biochemical Profiling of Structurally Validated Type II Inhibitors. J Med Chem 2015;58(1):466–479.

[31] Ung PMU, Rahman R, Schlessinger A. Redefining the Protein Kinase Conformational Space with Machine Learning. Cell Chem Biol 2018;25(7):916–924.e2.

[32] Möbitz H. The ABC of protein kinase conformations. BBA – Proteins and Proteomics 2015;1854(10, Part B):1555–1566.

[33] Gavrin LK, Saiah E. Approaches to discover non-ATP site kinase inhibitors. Med Chem Commun 2013;4:41–51.

[34] Arter C, Trask L, Ward S, Yeoh S, Bayliss R. Structural features of the protein kinase domain and targeted binding by small-molecule inhibitors. J Biol Chem 2022;298(8):102247.

[35] Wu P, Nielsen TE, Clausen MH. Small-molecule kinase inhibitors: an analysis of FDA-approved drugs. Drug Discov Today 2016;21(1):5–10.

[36] Fang Z, Grütter C, Rauh D. Strategies for the Selective Regulation of Kinases with Allosteric Modulators: Exploiting Exclusive Structural Features. ACS Chem Biol 2013;8(1):58–70.

[37] Wu P, Clausen MH, Nielsen TE. Allosteric small-molecule kinase inhibitors. Pharmacol Ther 2015;156:59–68.

[38] Wang B, Wu H, Hu C, Wang H, Liu J, Wang W, et al. An overview of kinase downregulators and recent advances in discovery approaches. Signal Transduct Target Ther 2021;6(1):423.

[39] Martinez R, Defnet A, Shapiro P. In: Shapiro P, editor. Avoiding or Co-Opting ATP Inhibition: Overview of Type III, IV, V, and VI Kinase Inhibitors Cham: Springer International Publishing; 2020. p. 29–59.

[40] Wright CJM, McCormack PL. Trametinib: First Global Approval. Drugs 2013;73(11):1245–1254.

[41] Henrik M, Wolfgang J, Cowan-Jacob SW. Expanding the Opportunities for Modulating Kinase Targets with Allosteric Approaches. Curr Top Med Chem 2017;17(1):59–70.

[42] Bhujbal SP, Jun J, Park H, Moon J, Min K, Hah JM. Gaining Insights into Key Structural Hotspots within the Allosteric Binding Pockets of Protein Kinases. Int J Mol Sci 2024;25(9):4725.

[43] Hu H, Laufkötter O, Miljković F, Bajorath J. Systematic comparison of competitive and allosteric kinase inhibitors reveals common structural characteristics. Eur J Med Chem 2021;214:113206.

[44] Laufkötter O, Hu H, Miljković F, Bajorath J. Structure- and Similarity-Based Survey of Allosteric Kinase Inhibitors, Activators, and Closely Related Compounds. J Med Chem 2022;65(2):922–934.

[45] Sturm N, Tinivella A, Rastelli G. Exploration and Comparison of the Geometrical and Physicochemical Properties of an α C Allosteric Pocket in the Structural Kinome. J Chem Inf Model 2018;58(5):1094–1103.

[46] Heinrich T, Gradler U, Bottcher H, Blaukat A, Shutes A. Allosteric IGF-1R inhibitors. ACS Med Chem Lett 2010;1(5):199– 203.

[47] Verma J, Vashisth H. Molecular basis for differential recognition of an allosteric inhibitor by receptor tyrosine kinases. Proteins 2024;92(8):905–922.

[48] Berendsen HJ, van der Spoel D, van Drunen R. GROMACS: A message-passing parallel molecular dynamics implementation. Comput Phys Commun 1995;91(1-3):43–56.

[49] Lindorff-Larsen K, Piana S, Palmo K, Maragakis P, Klepeis JL, Dror RO, et al. Improved side-chain torsion potentials for the Amber ff99SB protein force field. Proteins: Struct, Funct, Bioinf 2010;78(8):1950–1958.

[50] Jorgensen WL, Chandrasekhar J, Madura JD, Impey RW, Klein ML. Comparison of simple potential functions for simulating liquid water. J Chem Phys 1983;79(2):926–935.

[51] Chester C, Friedman B, Ursell F. An extension of the method of steepest descents. In: Mathematical Proceedings of the Cambridge Philosophical Society, vol. 53 Cambridge University Press; 1957. p. 599–611.

[52] Berendsen HJ, Postma Jv, Van Gunsteren WF, DiNola A, Haak JR. Molecular dynamics with coupling to an external bath. J Chem Phys 1984;81(8):3684–3690.

[53] Bussi G, Donadio D, Parrinello M. Canonical sampling through velocity rescaling. J Chem Phys 2007;126(1):014101.

[54] Schmidtke P, Bidon-Chanal A, Luque FJ, Barril X. MDpocket: open-source cavity detection and characterization on molecular dynamics trajectories. Bioinformatics 2011;27(23):3276–3285.

[55] Humphrey W, Dalke A, Schulten K. VMD: visual molecular dynamics. J Mol Graph 1996;14(1):33–38.

[56] Morris GM, Huey R, Lindstrom W, Sanner MF, Belew RK, Goodsell DS, et al. AutoDock4 and AutoDockTools4: Automated docking with selective receptor flexibility. J Comput Chem 2009;30(16):2785–2791.

[57] Hornak V, Abel R, Okur A, Strockbine B, Roitberg A, Simmerling C. Comparison of multiple Amber force fields and development of improved protein backbone parameters. Proteins 2006;65(3):712–725.

[58] Wang J, Wolf RM, Caldwell JW, Kollman PA, Case DA. Development and testing of a general amber force field. J Comput Chem 2004;25(9):1157–1174.

[59] Wang J, Wang W, Kollman PA, Case DA. Automatic atom type and bond type perception in molecular mechanical calculations. J Mol Graphics Modell 2006;25(2):247–260.

[60] Adasme MF, Linnemann KL, Bolz SN, Kaiser F, Salentin S, Haupt VJ, et al. PLIP 2021: expanding the scope of the protein–ligand interaction profiler to DNA and RNA. Nucleic Acids Res 2021 05;49(W1):W530–W534.

[61] Schrodinger L, The PyMOL molecular graphics system; 2015.

[62] David CC, Jacobs DJ. In: Livesay DR, editor. Principal Component Analysis: A Method for Determining the Essential Dynamics of Proteins Totowa, NJ: Humana Press; 2014. p. 193–226.

[63] Vashisth H, Maragliano L, Abrams CF. DFG-Flip in the Insulin Receptor Kinase Is Facilitated by a Helical Intermediate State of the Activation Loop. Biophys J 2012;102(8):1979–1987.

[64] Verma J, Vashisth H. Structural Models for a Series of Allosteric Inhibitors of IGF1R Kinase. Int J Mol Sci 2024;25(10):5368.

[65] Eswaran J, Knapp S. Insights into protein kinase regulation and inhibition by large scale structural comparison. BBA – Proteins and Proteomics 2010;1804(3):429–432.

[66] Rabiller M, Getlik M, Klüter S, Richters A, Tückmantel S, Simard JR, et al. Proteus in the World of Proteins: Conformational Changes in Protein Kinases. Archiv der Pharmazie 2010;343(4):193–206.

[67] Hobbs HT, Shah NH, Badroos JM, Gee CL, Marqusee S, Kuriyan J. Differences in the dynamics of the tandem-SH2 modules of the Syk and ZAP-70 tyrosine kinases. Protein Sci 2021;30(12):2373–2384.

[68] Taylor SS, Meharena HS, Kornev AP. Evolution of a dynamic molecular switch. IUBMB Life 2019;71(6):672–684.

[69] Goodwin NC, Cianchetta G, Burgoon HA, Healy J, Mabon R, Strobel ED, et al. Discovery of a Type III Inhibitor of LIM Kinase 2 That Binds in a DFG-Out Conformation. ACS Med Chem Lett 2015;6(1):53–57.

