## Supporting Information for "Conformational Dynamics in Insulin Receptor Kinase Reveals a Type III Allosteric Pocket"

### Table of Contents


**S1. Supplementary Methods** 3

Structure Selection and Preparation 3

Molecular Docking 3

Molecular Dynamics Simulations 4

Visualization & Analysis 5

**S2. Supplementary Figures** 7

**S3. Supplementary References** 12

### S1. Supplementary Methods

##### Structure Selection and Preparation

We used the tyrosine kinase domain of the human Insulin Receptor for our study. Among the available structures of the Insulin Receptor Kinase (IRK) resolved by X-ray crystallography in the Protein Data Bank, we selected the apo form of the IRK, representing its inactive conformation, which is not bound to ATP or any other molecules. The structure with the PDB ID 1IRK was selected for download. The missing atoms from the crystal structure were added using the Chimera interface of Modeller (Pettersen et al., 2004). This structure was used for further molecular dynamics (MD) simulations and molecular docking.

We used an experimentally known chemical compound, IUPAC name: N-1-[4-(5-cyano-1H-indol-3-yl)butyl]piperidin-4-yl-1H-indole-4-carboxamide, and performed molecular docking with the IRK to investigate its binding conformation. This inhibitor was selected based on its reported potency in inhibiting IRK activity ($IC_{50}$: 1.3 μM), as part of a series of indole butyl amine derivatives that have been experimentally validated for their inhibitory effects on IRK (Heinrich et al., 2010). Since no three-dimensional structure of the inhibitor was available in public databases, we generated its 3D coordinates using MarvinSketch 22.22 (ChemAxon). The structure was imported by entering the compound name as reported in the experimental study, *"1H-Indole-4-carboxylic acid{1-[4-(5-cyano-1H-indol-3-yl)-butyl]-piperidin-4-yl}-amide,"* into the MarvinSketch interface, and a 3D conformer was subsequently generated. The resulting coordinates of the ligand were saved in .mol2 format and used for subsequent docking studies.

##### Molecular Docking

Molecular docking was conducted using AutoDock 4.2 (Morris et al., 2009) using the IRK and inhibitor structures prepared above. Prior to docking, all crystallographic water molecules and heteroatoms (ethyl mercury ion) were removed manually using AutoDockTools (ADT), and polar hydrogen atoms were added to the IRK structure. Partial charges were applied to atoms to ensure appropriate electrostatic representation of the receptor. The structure was then saved in PDBQT format, the required format for AutoDock input, which contains atomic coordinates, partial charges, and AutoDock atom types.

For the ligand, we used a previously reported experimental inhibitor of IRK, for which 3D coordinates were generated as described above. The ligand .mol2 file was then imported into ADT, where Gasteiger charges were applied, and non-polar hydrogens were merged. Rotatable bonds were automatically detected, and the ligand was saved in PDBQT format.

The docking grid was centered on the predicted type III allosteric pocket of IRK. The grid center coordinates were as follows:

- X-dimension: 20.609 Å
- Y-dimension: 69.029 Å
- Z-dimension: 19.063 Å

The grid center was manually adjusted based on the known allosteric site region reported in our study. These parameters were saved in a grid parameter file (GPF) using AutoGrid4.

The docking parameter file (DPF) was configured with the following settings for robust sampling:

- Number of GA runs: 50
- Population size: 150
- Number of energy evaluations: 2500000
- Maximum number of generations: 27000
- Elitism: 1
- Mutation rate: 0.02
- Crossover rate: 0.8
- Local search frequency: 0.06

Docking was performed using the Lamarckian Genetic Algorithm (Morris et al., 1998) implemented in AutoDock4. The docked poses were ranked by predicted binding energy, and the lowest-energy conformation was selected for subsequent molecular dynamics simulations. The coordinates of the docked protein and the ligand were saved as a complex in the PDB format. They were also saved individually for topology generation in the MD simulation protocol.

**Molecular Dynamics (MD) Simulations**

Molecular dynamics simulations were performed using GROMACS 2020.4 (Berendsen et al., 1995) for the apo IRK and the inhibitor bound IRK (obtained from docking) structures. To prepare each system for production MD simulations, a standard protocol of system solvation, neutralization, minimization and equilibration was followed. Prior to this protocol, the protein and ligand structures were prepared to generate GROMACS compatible topology.

Topology Generation (Protein): The `pdb2gmx` module was run on the protein structure of each system (apo and docked protein) for assigning appropriate force field parameters and generating the required topology and coordinate files for GROMACS simulations. The force field used for the protein atoms was AMBER ff999SB-ILDN (Lindorff-Larsen et al., 2010).

Topology Generation (Ligand): The inhibitor structure parameters were generated using the Antechamber module of AmberTools. The atom types were based on GAFF, which is a general force field for organic molecules and AM1-BCC charges were applied. Parmchk and tleap were used to generate Amber compatible topology files. The conversion to AMBER topology (.prmtop) and coordinate (.inpcrd) files to GROMACS compatible topology (.top) and coordinate (.gro) formats was performed using the ParmEd library in python.

Solvation and Neutralization: The system (apo IRK or IRK+Inhibitor) was solvated in a cubic simulation domain with TIP3P explicit water molecules (Jorgensen et al., 1983), ensuring a minimum of 10 Å from the box edge to any atom of the protein. Sodium ($Na^+$) and chloride ($Cl^-$) ions were added to neutralize the system by passing the flag "-neutral" in the genion command. The total number of atoms in each system was as follows: apo IRK – 56,584 atoms, and IRK with inhibitor – 56,598 atoms.

For all dynamics steps, the bond parameters, non-bonded and electrostatic settings were as follows:

The LINCS algorithm (B. Hess et al., 1997) was used to apply holonomic constraints on all bonds involving hydrogen atoms, with lincs order = 4 and iteration = 1 to ensure constraint accuracy. For non-bonded interactions, Verlet cutoff scheme was used with a neighbor search frequency of every 10 steps using the grid-based method. The short-range electrostatic and van der Waals interactions cutoff were 1.0 nm. For electrostatics, the Particle Mesh Ewald (PME) (U. Essmann et al., 1995) method was used with a fourth-order interpolation (pme_order = 4) and a Fourier grid spacing of 0.16 nm to accurately treat long-range Coulomb interactions. The time-step used in all simulations was 2 femtoseconds. Periodic boundary conditions were applied in all directions.

<u>Minimization</u>: Energy minimization was performed using steepest descent algorithm (Chester et al., 1957). A maximum of 50,000 minimization steps were performed with a step size (emstep) of 0.01 nm, and a convergence criterion (emtol) of force < 1000 kJ $mol^{-1}$ $nm^{-1}$.

<u>Equilibration Phase 1 (NVT ensemble):</u> In the first equilibration step, the system was simulated in the NVT ensemble, maintaining a constant number of particles, volume, and temperature at 300 K for 500 ps. Temperature control was achieved using the velocity-rescale thermostat (G. Bussi et al., 2007) with a coupling constant (tau_t) of 0.1 ps. To prevent large deviations in the protein structure, positional restraints were applied to the protein heavy atoms and, if present (in IRK-Inhibitor system), to the ligand heavy atoms. The force-constant for these restraints was set to 1000 kJ/mol/$nm^2$.

<u>Equilibration Phase 2 (NPT ensemble):</u> In the second equilibration step, the system was simulated in the NPT ensemble, maintaining a constant number of particles, pressure, and temperature at 300 K and 1 bar for 10 ns. Temperature was maintained using the velocity-rescale thermostat (tau_t = 0.1 ps), and pressure was controlled using the Berendsen barostat (Berendsen et al., 1984) with a coupling constant (tau_p) of 2.0 ps and compressibility of 4.5e-5 $bar^{-1}$. Positional restraints were retained as in the NVT ensemble.

<u>Production MD:</u> In the production MD step, three independent NPT ensemble simulations were conducted for each system (apo IRK and IRK+Inhibitor), each for 1 μs with a timestep of 2 fs, totaling 3 μs of simulation time for each system. Trajectory outputs (coordinates, velocities, and energies) were saved every 100 ps. Further, gromacs trjconv was used to post-process each trajectory for analysis.

**Visualization & Analysis**

Trajectories were analyzed and visualized using GROMACS modules (version 2020.4) and VMD (version 1.9.3). All the plots were generated using MATLAB (R2024a) (The MathWorks Inc. 2024).

1. <u>RMSD:</u> We used GROMACS module rms to calculate the root mean squared deviation of the protein backbone atoms. The initial frame of the simulation trajectory (equilibrated) was used as a reference. GROMACS make_ndx was used to extract the atoms corresponding to the αC-helix, A-loop and GC-loop for calculations. To evaluate RMSD for the inhibitors, heavy atoms of the inhibitor considered taking the initial docked conformation as a reference.

2. <u>RMSF:</u> GROMACS module rmsf was used for the calculation. The root mean squared fluctuation for Cα atoms of each residue was calculated from their mean positions.

3. <u>Principal Component Analysis (PCA):</u> PCA was performed using GROMACS tools to examine conformational transitions in apo-IRK and inhibitor-bound IRK. A covariance matrix of backbone atomic fluctuations was generated using gmx covar, and eigenvectors/eigenvalues were obtained by diagonalization. The most significant motions were identified from the top principal components (PCs). Trajectories were projected onto PC1 and PC2 using gmx anaeig to define the essential subspace. The free energy landscape was computed using gmx sham, projecting the data along PC1 and PC2 to identify lowest-energy conformational states.

4. <u>Probability Density Distribution:</u> We used the ksdensity function in MATLAB to evaluate the probability density estimate of the residue interaction distances.

5. <u>Protein-Ligand Interaction:</u> PLIP web tool (Link: https://plip-tool.biotec.tu-dresden.de) was used to evaluate IRK-inhibitor interactions and visualized with PyMol.

#### S2. Supplementary Figures

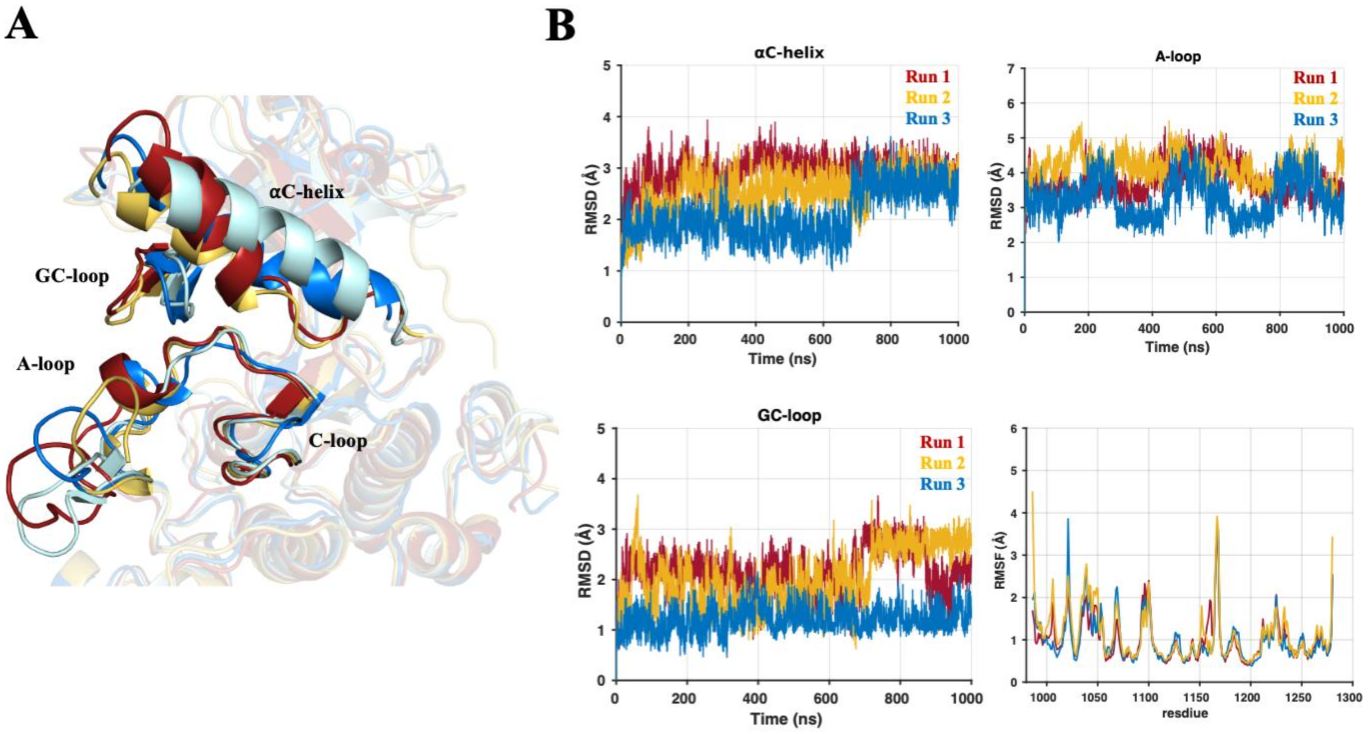


Figure S1: (A) Structural superimposition of the apo IRK conformations. IRK conformations: X-ray (Cyan); Run 1 (Red); Run 2 (Yellow); Run 3 (Blue). (B) (upper-left panel) root mean squared deviation (RMSD) of the backbone atoms of the αC-helix, (upper-right panel) RMSD of the backbone atoms of A-loop, (lower-left panel) RMSD of the backbone atoms of the GC-loop and, (lower-right panel) root mean squared fluctuation (RMSF) of the Cα atoms.

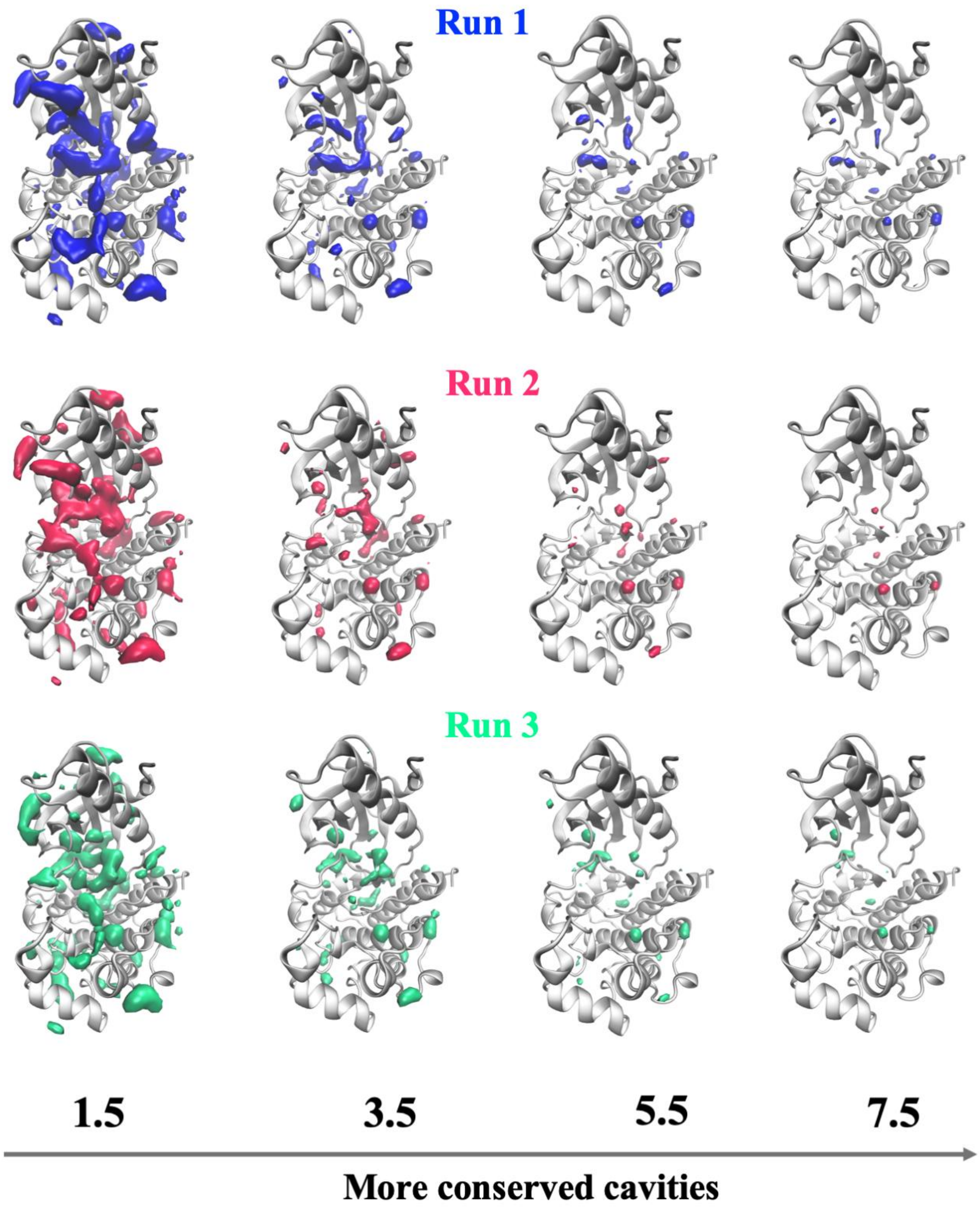


Figure S2: Density grid maps obtained from each of the three independent MD simulation trajectories of apo IRK. The grid maps shown have different threshold of isovalues as labeled for each column.

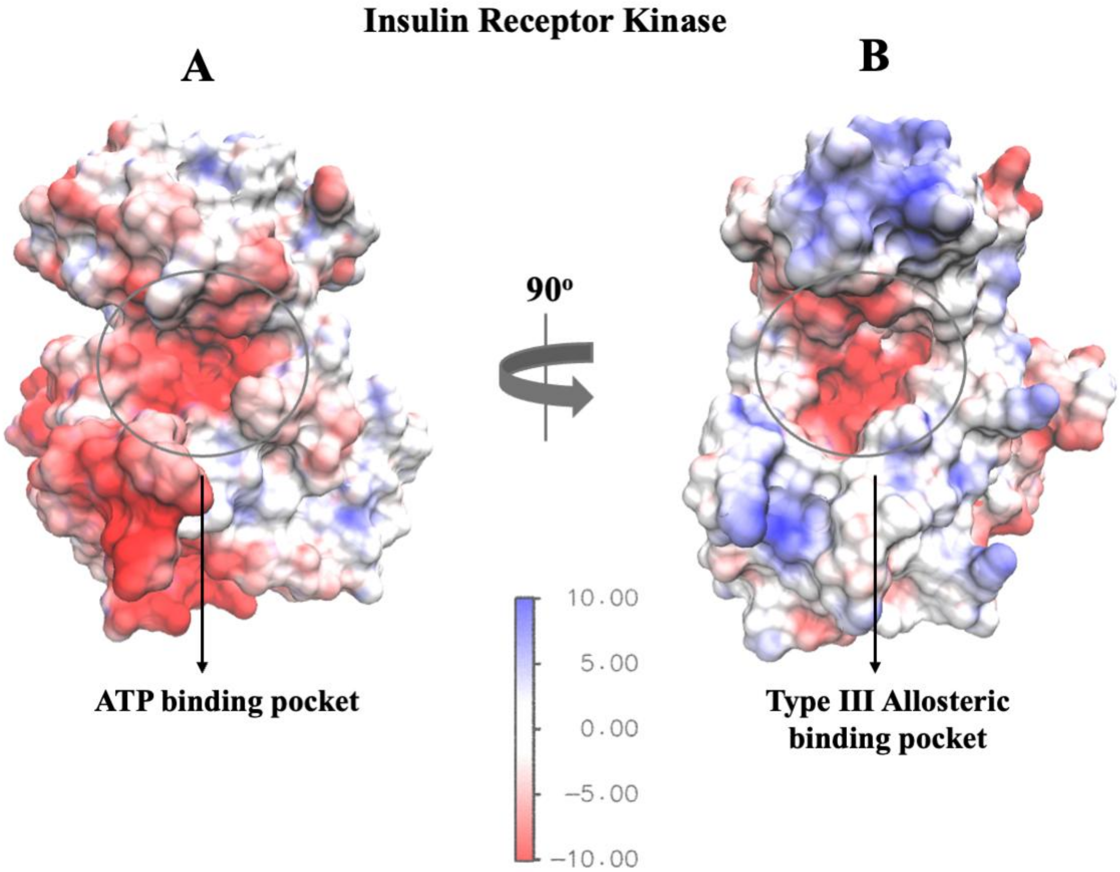


Figure S3: Electrostatic potential energy surface map for the IRK domain in its inactive conformation (PDB ID: 1IRK). (A) A surface view showing the ATP binding pocket region. (B) A surface view showing the type III allosteric pocket region.

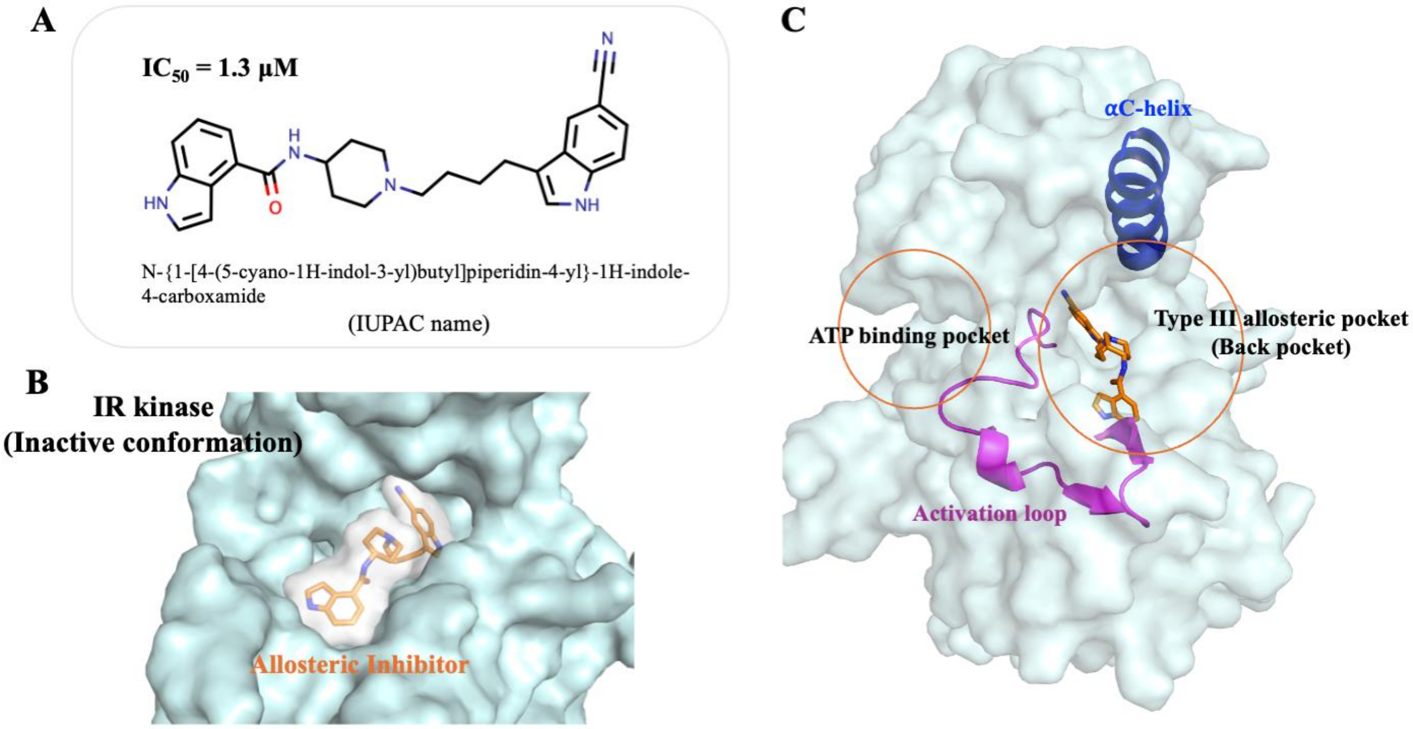


Figure S4: (A) Chemical structure of the allosteric inhibitor of IRK studied in our work. (B) The docked conformation of the inhibitor in the allosteric pocket of the IRK domain. (C) Structure of the IRK domain with the docked conformation of the inhibitor positioned within the type III allosteric pocket, also known as the “back pocket”, in relation to the ATP pocket (orthosteric site).

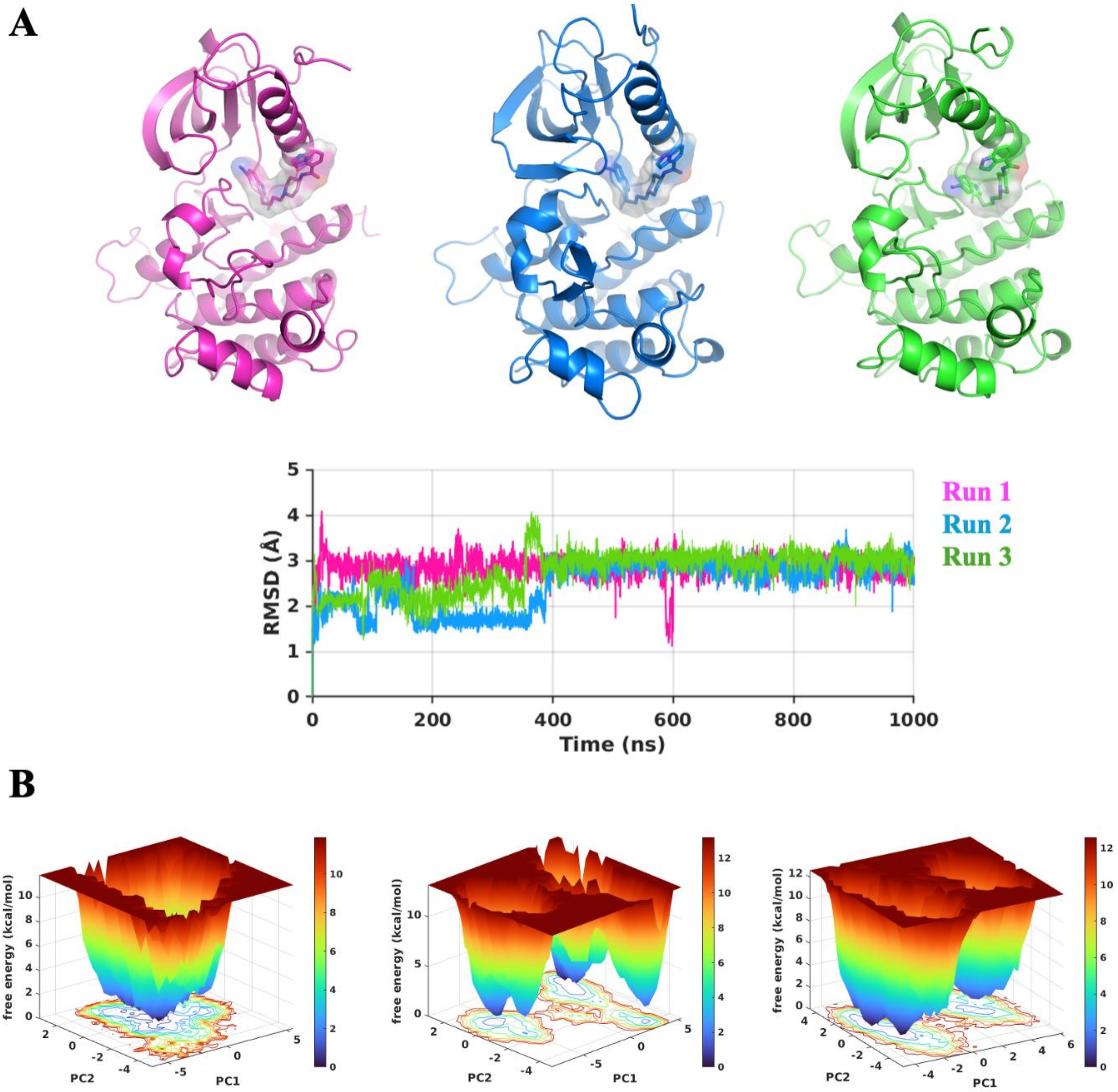


Figure S5: (A) Root mean squared deviation (RMSD) of the protein backbone atoms for the inhibitor-bound IRK. IRK-inhibitor bound conformations obtained from the FES are also shown. (B) Projections of the FES along two principal components (PC1 and PC2) for the IRK-inhibitor bound complex.

#### S2. Supplementary References


B. Hess, H. Bekker, H.J.C. Berendsen, and J.G.E.M. Fraaije. (1997) LINCS: A Linear Constraint Solver for Molecular Simulations. J. Comp. Chem., 18, 1463–72.

Berendsen, H. J.C., J. P.M. Postma, W. F. Van Gunsteren, A. Dinola, and J. R. Haak. (1984) Molecular Dynamics with Coupling to an External Bath. J. Chem. Phys., 81, 3684–90.

Berendsen, H. J.C., D. van der Spoel, and R. van Drunen. (1995) GROMACS: A Message-Passing Parallel Molecular Dynamics Implementation. Comput. Phys. Commun., 91, 43–56.

Chester, C., B. Friedman, and F. Ursell. (1957) An Extension of the Method of Steepest Descents. Mathematical Proceedings of the Cambridge Philosophical Society, 53, 599–611.

G. Bussi, D. Donadio, and M. Parrinello. (2007) Canonical Sampling through Velocity Rescaling. J. Chem. Phys., 126, 014101.

Heinrich, Timo, Ulrich Grädler, Henning Böttcher, Andree Blaukat, and Adam Shutes. (2010) Allosteric IGF-1R Inhibitors. ACS Med. Chem. Lett., 1, 199–203.

Lindorff-Larsen, Kresten and Piana, Stefano and Palmo, Kim and Maragakis, Paul and Klepeis, John L. and Dror, Ron O. and Shaw, David E. (2010) Improved Side-Chain Torsion Potentials for the Amber Ff99SB Protein Force Field. Proteins, 78, 1950–58.

Morris, G. M., Goodsell, D. S., Halliday, R.S., Huey, R., Hart, W. E., Belew, R. K. and Olson, A. (1998) Automated Docking Using a Lamarckian Genetic Algorithm and and Empirical Binding Free Energy Function. J. Com. Chem., 19, 1639–62.

Morris, Garrett M., Huey Ruth, William Lindstrom, Michel F. Sanner, Richard K. Belew, David S. Goodsell, and Arthur J. Olson. (2009) AutoDock4 and AutoDockTools4: Automated Docking with Selective Receptor Flexibility. J. Com. Chem., 30, 2785–91.

Pettersen, Eric F., Thomas D. Goddard, Conrad C. Huang, Gregory S. Couch, Daniel M. Greenblatt, Elaine C. Meng, and Thomas E. Ferrin. (2004) UCSF Chimera--a Visualization System for Exploratory Research and Analysis. J. Comp. Chem., 25, 1605–12.

The MathWorks Inc. (2024). MATLAB Version: (R2024a). Natick, Massachusetts: The MAthWorks Inc.

U. Essmann, L. Perera, M.L. Berkowitz, T. Darden, H. Lee, and L.G. Pedersen. (1995) A Smooth Particle Mesh Ewald Potential. J. Chem. Phys., 103, 8577–92.
